# Neurochemical and Neurophysiological Effects of Intravenous Administration of *N,N*-Dimethyltryptamine in Rats

**DOI:** 10.1101/2024.04.19.589047

**Authors:** Nicolas G. Glynos, Emma R. Huels, Amanda Nelson, Youngsoo Kim, Robert T. Kennedy, George A. Mashour, Dinesh Pal

## Abstract

*N*,*N*-dimethyltryptamine (DMT) is a serotonergic psychedelic that is being investigated clinically for the treatment of psychiatric disorders. Although the neurophysiological effects of DMT in humans are well-characterized, similar studies in animal models as well as data on the neurochemical effects of DMT are generally lacking, which are critical for mechanistic understanding. In the current study, we combined behavioral analysis, high-density (32-channel) electroencephalography, and ultra-high-performance liquid chromatography-tandem mass spectrometry to simultaneously quantify changes in behavior, cortical neural dynamics, and levels of 17 neurochemicals in medial prefrontal and somatosensory cortices before, during, and after intravenous administration of three different doses of DMT (0.75 mg/kg, 3.75 mg/kg, 7.5 mg/kg) in male and female adult rats. All three doses of DMT produced head twitch response with most twitches observed after the low dose. DMT caused dose-dependent increases in serotonin and dopamine levels in both cortical sites along with a reduction in EEG spectral power in theta (4-10 Hz) and low gamma (25-55 Hz), and increase in power in delta (1-4 Hz), medium gamma (65-115 ), and high gamma (125-155 Hz) bands. Functional connectivity decreased in the delta band and increased across the gamma bands. In addition, we provide the first measurements of endogenous DMT in these cortical sites at levels comparable to serotonin and dopamine, which together with a previous study in occipital cortex, suggests a physiological role for endogenous DMT. This study represents one of the most comprehensive characterizations of psychedelic drug action in rats and the first to be conducted with DMT.

**Significance Statement:** *N*,*N*-dimethyltryptamine (DMT) is a serotonergic psychedelic with potential as a tool for probing the neurobiology of consciousness and as a therapeutic agent for psychiatric disorders. However, the neurochemical and neurophysiological effects of DMT in rat, a preferred animal model for mechanistic studies, are unclear. We demonstrate that intravenous DMT caused a dose-dependent increase in serotonin and dopamine in medial prefrontal and somatosensory cortices, and simultaneously increased gamma functional connectivity. Similar effects have been shown for other serotonergic and atypical psychedelics, suggesting a shared mechanism of drug action. Additionally, we report DMT during normal wakefulness in two spatially and functionally distinct cortical sites — prefrontal, somatosensory — at levels comparable to those of serotonin and dopamine, supporting a physiological role for endogenous DMT.

## Introduction

Serotonergic psychedelics, including *N*,*N*-dimethyltryptamine (DMT), produce profound alterations in perception, cognition, mood, and affect in humans (Nichols, 2016) and have been shown to have therapeutic potential for treating a variety of psychiatric disorders (Palhano-Fontes et al., 2019; D’Souza et al., 2022; Goodwin et al., 2022; Holze et al., 2022). DMT has been reported in human blood, urine, and cerebrospinal fluid (Barker et al., 2012) and a recent *in vivo* study revealed the presence of endogenous DMT in occipital cortex, including pineal gland (Dean et al., 2019). However, whether the endogenous DMT is solely restricted to occipital cortex and pineal gland or is present in other cortical areas, is not known. Of note, in contrast to other serotonergic psychedelics, the consciousness-altering effects of DMT are relatively short in duration. When administered intravenously to humans, peak effects occur within approximately three minutes of administration and persist for less than fifteen minutes (Timmermann et al., 2023). The profound and short-acting effects of DMT make it a valuable tool for probing the neurobiology of consciousness, and for future psychedelic-assisted therapies compared to psilocybin or lysergic acid diethylamide (LSD), where dosing sessions can last up to 10 hours (Nichols, 2016). Additionally, the study of endogenous DMT has the potential to provide insights into the mechanisms of action of other serotonergic psychedelics, via an investigation of its properties as a potential neurotransmitter that mediates consciousness or other functions (Vargas et al., 2023).

It has been established that activation of serotonin-2A (5-HT_2A_) receptors plays a role in mediating the psychoactive effects of DMT (Keiser et al., 2009), but less is known about the neurochemical changes resulting from DMT administration. Early rat studies, in which post-mortem concentrations of neurotransmitters or their metabolites were analyzed after intraperitoneal administration of DMT, showed increases in extracellular dopamine and its primary metabolites (Haubrich and Wang, 1977; Smith, 1977; Waldmeier and Maître, 1977). Serotonin levels have also been reported to increase in a dose dependent manner following intraperitoneal administration of DMT in rats (Freedman et al., 1970). However, these studies relied on post-mortem analysis of tissue homogenates, lacked spatial resolution, and utilized coarse approaches (e.g., fluorometric analyses), as compared to the more technically sophisticated and robust methodologies in current use (e.g., ultra-high-performance liquid chromatography-tandem mass spectrometry).

Recent human studies using intravenous and inhaled administration of DMT (Timmermann et al., 2019, 2023; Pallavicini et al., 2021) showed a significant reduction in EEG spectral power in alpha and beta bands, along with simultaneous increase in spectral power in delta and gamma range. Additionally, increased corticocortical coherence and signal diversity have also been demonstrated in the gamma range following DMT administration. However, there have been no studies to characterize the effects of DMT administration on neurophysiological dynamics in rodents, which are a valuable translational model for mechanistic studies.

Therefore, in the current study, we determined the effect of intravenous administration of DMT on changes in 1) behavior, 2) levels of 17 neurochemicals including acetylcholine, serotonin, dopamine, γ-aminobutyric acid, and glutamate in medial prefrontal cortex and somatosensory cortex, and 3) EEG spectral power and functional connectivity across the cortex in rats.

## Materials and Methods

### Rats

The experiments were approved by the Institutional Animal Care and Use Committee at the University of Michigan and were performed in accordance with the Guide for the Care and Use of Laboratory Animals (8^th^ Edition, The National Academies Press), as well as ARRIVE guidelines. Adult male and female Sprague Dawley rats (n = 19, 9 male and 8 female, 300-500g, Charles River Laboratories) were used for all experiments and were housed in a temperature- and light-controlled facility (12 hours light:12 hours dark cycle, lights on at 8:00 AM), with ad libitum access to food and water.

### Surgical procedures

Rats were anesthetized in an induction chamber using 4-5% isoflurane in 100% oxygen. After anesthetic induction, the head was shaved and the rats were immobilized in a stereotaxic frame using blunt ear bars (Model 963, David Kopf Instruments, Tujanga, CA). The rats were positioned to breathe isoflurane through a nose cone (Model 906, David Kopf Instruments, Tujanga, CA), which was titrated to maintain loss of the pedal and palpebral reflex. The delivered isoflurane concentration was monitored throughout the surgical procedure using an anesthetic agent analyzer (Capnomac Ultima, Datex Medical Instrumentation, Tewksbury, MA). Body temperature was maintained at 37°C using a far-infrared heating pad (RT-0502, Kent Scientific, Torrington, CT). Before surgery, subcutaneous buprenorphine (0.01 mg/kg; Buprenex, Par Pharmaceutical, Chestnut Ridge, NY; NDC 42023-179-05) and carprofen (5 mg/kg; Zoetis; NADA #141-199) were administered as presurgical analgesics, and cefazolin (20 mg/kg; West-Ward-Pharmaceutical, Eatontown, NJ; NDC 0143-9924-90) was administered as a prophylactic antibiotic. A sagittal scalp incision was made, and the connective tissue was cleared to expose the skull. Thirty burr holes were drilled across the skull for the implantation of custom-made stainless-steel EEG screw electrodes in a regularly spaced grid of 8 rows that was parallel to coronal suture and spanned the area between 4 mm anterior and 10 mm posterior to bregma; across the 8 rows, the electrodes were arranged in columns parallel and lateral (2 mm to 4.5 mm) to the mid-sagittal suture. Screw electrodes were implanted above the nasal sinus and cerebellum to serve as the reference and ground electrodes, respectively. Two concentric open flow microperfusion (OFM) guide tubes (cOFM-GD-8-2, BASi Research Products, West Lafayette, IN) with dummy inserts (cOFM-D-10 and cOFM-Lock) were implanted through burr holes aimed at medial prefrontal cortex (mPFC) (from Bregma: 3.0 mm anterior, 0.5 mm mediolateral, and 3.0 mm ventral) and somatosensory barrel field cortex (S1BF) (from Bregma: 3.24 mm posterior, 5.3 mm mediolateral, and 4.0 mm ventral) (Paxinos and Watson, 2007). In addition, all rats were surgically fitted with an indwelling catheter (MRE-040, Micro-Renathane tubing, Braintree Scientific, Braintree, MA) in the jugular vein for intravenous infusion of DMT fumarate (Cayman Chemical, Ann Arbor, MI). All electrodes were secured to a 32-pin Mill-Max connector (Mouser Electronics, Mansfield, TX) and the entire assembly was secured to the cranium with dental cement. Post-surgical buprenorphine (0.03 mg/kg) was administered subcutaneously every 8-12 hours for 48 hours to provide analgesia. The rats were allowed a minimum of 14 days of post-surgical recovery.

### Histological analysis

At the completion of experiments, the rats were euthanized with carbon dioxide inhalation and intracardially perfused with 150 mL of wash solution containing 0.1 M (pH 7.2) phosphate buffered saline (1219SK, EM Sciences, Hatfield, PA) and 1% heparin (1 unitl/mL, Sagent Pharmaceuticals, Schaumburg, IL) followed by 200 mL of a fixative solution containing 4% paraformaldehyde and 4% sucrose in 0.1 M phosphate buffer (pH 7.2). Brains were extracted and stored in the fixative solution for at least 24 hours at 4°C, and then transferred to a 30% sucrose in phosphate buffer solution. Coronal sections (30 µm) were cut through the mPFC and S1BF with a cryostat (Leica Microsystems, Nussloch, Germany). The brain sections were mounted on glass slides and stained with Cresyl violet to confirm the site of OFM probe insertion in mPFC and S1BF regions.

### Experimental design

The experimental design is illustrated in **figure 1A**. The EEG electrode montage and the approximate location of OFM probes is illustrated in **figure 1B**. The rats were conditioned to the experimental chambers and EEG recording cables for at least one week before starting the experiments. On the day of the experiment, rats were connected to the EEG recording cable and OFM probes (cOFM-S-10, BASi Research Products, West Lafayette, IN) were lowered into mPFC and S1BF. The probes were connected to a peristaltic microperfusion pump (MPP102 PC, Joanneum Research, Graz, Austria) for continuous perfusion (1 µL/min) with artificial cerebrospinal fluid (aCSF; 145 mM NaCl, Sigma Aldrich S9888; 2.68 mM KCl, Sigma Aldrich P9333; 1.40 mM CaCl_2_•2H_2_O, Sigma Aldrich C8106; 1.01 mM MgSO_4_•7H_2_O, Sigma Aldrich 63140; 1.55 mM Na_2_HPO_4_, Sigma Aldrich S5136; 0.45 mM NaH_2_PO_4_•H2O, Sigma Aldrich 52074; pH 7.4). Data collection began at least one hour after connecting the rats to the EEG cables and OFM probes. EEG and OFM data were collected simultaneously and continuously for the duration of the experiment, which lasted 137.5 minutes. The OFM samples were collected in 12.5 min epochs (1 µL/min) simultaneously from both mPFC and S1BF regions and consisted of 4 baseline epochs (B1-B4) before the intravenous administration of DMT followed by 7 drug epochs (D1-D7) during and after the intravenous administration of DMT. To keep the behavioral state constant during pre-DMT baseline condition, we tapped on the recording chamber if the rat assumed a sleeping posture and/or if slow waves, indicative of drowsiness, appeared in the EEG. At the start of the 5^th^ epoch (i.e., D1 after baseline control), DMT-fumarate dissolved in saline was administered over a 5-minute period (100 µL/min) as an intravenous bolus at one of the following doses: 1) low - 0.75 mg/kg, 2) medium - 3.75 mg/kg, and 3) high - 7.5 mg/kg in a random order with at least 5 days between each dose. Thus, in this within-subject repeated-measures design, each rat served as a control for itself, and each experimental session started with the collection of four 12.5-minutes long baseline data epochs (behavioral, neurochemical, and EEG) before the intravenous administration of DMT. The EEG data were recorded from 17 (9 male) rats for each dose. For the neurochemical experiments, out of the total 57 experimental sessions (19 rats x 3 doses), the mPFC probes failed during 5 experiments and S1BF probes failed during 7 experiments. The final sample sizes for neurochemical data are as follows: low dose mPFC: 16 (8 males); low dose S1BF: 14 (8 males); medium dose mPFC: 16 (9 males); medium dose S1BF: 15 (8 males); high dose mPFC: 14 (7 males); high dose S1BF: 15 (8 males). The OFM samples were collected in a refrigerated fraction collector (4°C) and immediately processed and frozen (−80°C) for ultra-high-performance liquid chromatography-tandem mass spectrometry (uHPLC-MS/MS) analysis. All experiments were video recorded for later behavioral analysis by an investigator blinded to the experimental condition. At the conclusion of data collection, the OFM sample collection sites were histologically verified (Fig. 1C-D).

**Figure 1.**
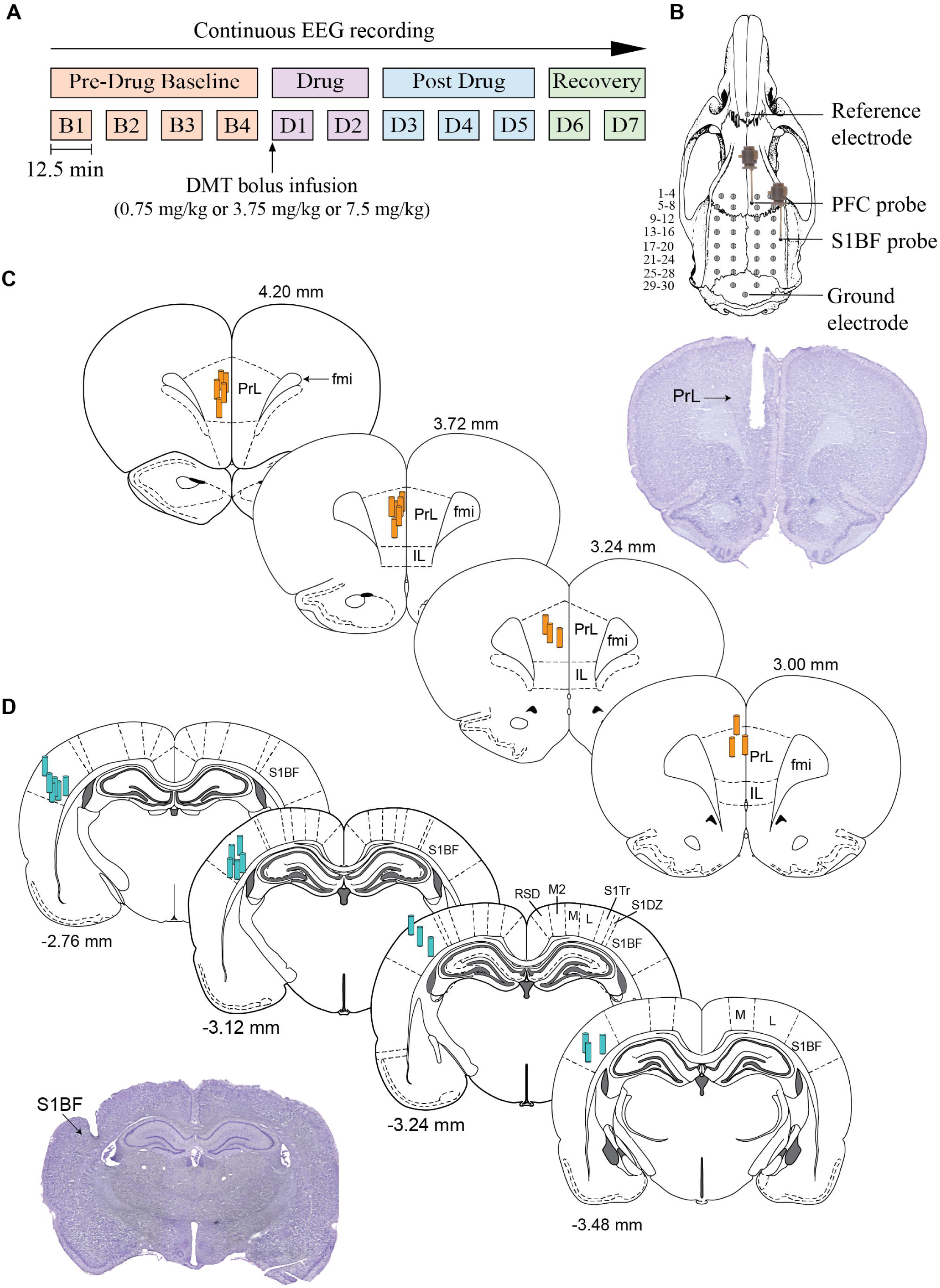
Schematic showing the experimental timeline and histological verification of the neurochemical sampling sites. Panel (A) shows the experimental design and timeline. A perfusate sample was collected from both medial prefrontal cortex (mPFC) and somatosensory cortex (S1BF) for each 12.5 min epoch before (pre-drug baseline condition) and after DMT administration. DMT was administered at the start of the D1 epoch over a 5-minute period (100 µL/min) as an intravenous bolus at one of the following doses: 1) low - 0.75 mg/kg, 2) medium - 3.75 mg/kg, and 3) high - 7.5 mg/kg in a random order with at least 5 days between each dose. The perfusate samples were prepared for neurochemical quantification via ultra-high-performance liquid chromatography-tandem mass spectrometry (uHPLC-MS/MS) as described in the methods section. See Extended Data Table 1-1 for a list of analytes and multiple reaction monitoring scanning parameters used in uHPLC-MS/MS analyses. EEG was collected continuously during the entire experimental session. Panel (B) shows the EEG montage (30 screw electrodes) to record high-density intracranial EEG and placement of open flow microperfusion (OFM) neurochemical sampling probes in mPFC and S1BF. Electrodes are numbered 1-30, row-wise, starting with the upper left electrode. Screw electrodes implanted above the sinus and cerebellum served as reference and ground electrodes, respectively. Schematics from the rat brain atlas (Paxinos and Watson, 2007) show coronal sections through mPFC (C) and S1BF (D). Orange and turquoise cylinders represent the sampling length (1 mm) of the microperfusion probes implanted in mPFC and S1BF, respectively. The numbers associated with each stereotaxic map indicate the distance relative to bregma; positive numbers indicate anterior to bregma, and negative numbers indicate posterior to bregma. Representative cresyl violet-stained coronal brain sections (30 µm) are shown for the histological verification of the probe placement in mPFC (C) and S1BF (D). fmi: forceps minor corpus callosum, IL: infralimbic cortex, L: lateral parietal association cortex, M: medial parietal association cortex, M2: secondary motor cortex, PrL: prelimbic region, RSD: retrosplenial dysgranular cortex, S1DZ: primary somatosensory cortex dysgranular zone, S1Tr: primary somatosensory trunk region.

### Behavioral analysis

All experiments were video recorded and data collection included 17 rats for each of the low and high doses, and 15 rats for the medium dose. The details of the experimental sessions were masked to allow for blinded analysis. General behavioral patterns were observed and documented for the duration of each experiment. Head twitch response (HTR), defined as a rapid, side-to-side rotational movement of the head independent of grooming behavior, was quantified during a 15-minute window, starting at the beginning of the DMT infusion (D1).

### Neurochemical quantification via ultra-high-performance liquid chromatography-tandem mass spectrometry (uHPLC-MS/MS)

To prepare perfusate samples for uHPLC-MS/MS analysis, 10 µL of sample was mixed with 2.5 µL of carbonate buffer (200 mM), and derivatized with 25 µL of 2% benzoyl chloride (Sigma Aldrich 259950) in acetonitrile (v/v), and the mixture was centrifuged for 10 minutes to precipitate proteins. To normalize for extraction efficiency and mass spectrometry ionization efficiency, 9 µL of the supernatant was spiked with 11 µL of internal standard, which contained ^13^C- or deuterium-labeled derivatives of all analytes of interest (see **Extended Data Table 1-1** for a list of analytes and multiple reaction monitoring scanning parameters). Prior to each uHPLC-MS/MS analysis, 6-point calibration curves were constructed for each analyte based on the peak area ratio of the calibration standard to the internal standard by linear regression. Calibration standards consisted of a mixture of all compounds of interest and were spiked with the corresponding ^13^C- or deuterium-labeled internal standard. All samples and standards were analyzed using a Phenomenex Kinetex C18 chromatography column (100 x 2.1 mm, 1.7 µm, 100 Å) on a Vanquish uHPLC (Thermo Fisher Scientific, Gemering, Germany) interfaced to a TSQ Quantum Ultra triple quadrupole mass spectrometer (Thermo Fisher Scientific, San Jose, CA). Mobile phase A was 10 mM ammonium formate with 0.15% (v/v) formic acid in water. Mobile phase B was acetonitrile. The gradient used was as follows: initial, 5% B; 0.01 min, 19% B; 0.68 min, 26% B; 1.05 min, 75% B; 1.8 min, 100% B; 2.2 min, 100% B; 2.3 min, 5% B; 3.0 min, 5% B. The flow rate was maintained at 600 μL/min and the sample injection volume was 7.5 µL. The autosampler was kept at ambient temperature, with the column being held at 30°C in still air mode. Electrospray ionization was used in positive mode at 4 kV. The capillary temperature was 325°C, the vaporizer temperature was 300°C, the sheath gas pressure was 50 Arb, and the auxiliary gas pressure was 10 Arb. Ions were detected in tandem mass spectrometry mode. Automatic peak integration was performed using XCalibur 3.0 MS software, and all peaks were visually inspected to ensure accurate integration. The following 17 analytes were included in neurochemical analyses: acetylcholine, adenosine, aspartate, choline, dopamine, DMT, GABA, glucose, glutamate, glutamine, glycine, histamine, homovanillic acid, phenylalanine, serine, serotonin, and taurine.

### EEG data acquisition and analysis

Monopolar EEG data were acquired using a 32-channel head stage (Cereplex μ, Blackrock Microsystems, Salt Lake City, UT). The signals were digitized at 1 kHz and bandpass filtered between 0.1-500 Hz using a Cereplex Direct system paired with the Cereplex Direct Software Suite (Blackrock Microsystems, Salt Lake City, UT). Raw EEG signals were imported into MATLAB (version 2022b; MathWorks, Inc.; Natick, MA) and data segments with movement or hardware artifacts were removed after identification by visual inspection of both the raw waveform and spectrogram of the EEG signals in 10-second windows. EEG segments containing slow-waves, indicative of sleep or a sleep-like state, were excluded. Bad channels were also visually identified and excluded. Segments of clean EEG data (total lengths between 1 and 5 minutes per epoch) were subsequently used to compute power spectrum density (PSD) and weighted phase lag index (wPLI) within each of the following frequency bands: delta (1-4 Hz), theta (4-10 Hz), low gamma (25-55 Hz), medium gamma (65-115 Hz), and high gamma (125-155 Hz). PSD was calculated using Matlab’s implementation of Welch’s method (pwelch.m) at the channel-level on clean data segments (10-second moving window, 90% overlap, 2-second Hamming window) and then averaged across all channels. Relative power was calculated as the absolute power within a given frequency divided by the total absolute power across all frequencies for each of the 11 epochs. wPLI is a measure of functional connectivity based on the phase relationship between two signals and is robust to volume conduction (Vinck et al., 2011). Before computing wPLI, the full continuous data recordings were bandpass filtered to the desired frequency band using FieldTrip’s implementation of a windowed-sinc finite impulse response filter (Oostenveld et al., 2010). Artifact-free segments (between 1 and 5 minutes) of the filtered EEG data — identical to those used for PSD — were extracted for analysis. A Hilbert transform was used to extract the analytic signal from each electrode. The complex conjugate of each channel pair (***xy***) was subsequently used to estimate the cross spectrum (*c*_*xy*_), with the imaginary component of the cross spectrum being 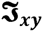 wPLI was then estimated for 10-second non-overlapping windows using the following equation:

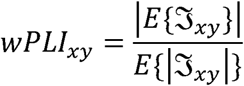

wPLI values were subsequently normalized using surrogate data (n = 50) generated through phase-shuffling while preserving the amplitude distribution of each signal. The normalized wPLI (*wPLJ*_*norm*_) was then computed using the mean (*µ*_*shuff*_) and standard deviation (*σ*_*shuff*_ ) of the surrogate data:

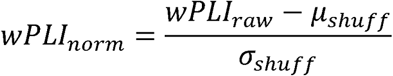

Global wPLI was computed by averaging across all channel pairs for each frequency band for each of the 11 epochs.

### Statistical analysis

All statistical analyses were completed with R software (RStudio version 2023.09.1+494). The sample size was decided based on a priori power analyses conducted using neurochemical and neurophysiological pilot data from DMT studies in our lab and was informed by the neurochemical data (changes in acetylcholine) from a previous publication (Brito et al., 2021). Primary outcome measures included DMT-induced HTR, changes in the levels of neurotransmitters, and the changes in EEG measures of relative spectral power and functional connectivity (wPLI). For analysis of HTR data, a linear mixed model was implemented, with ‘dose’ treated as a fixed factor and ‘subject’ treated as a random intercept. An alpha threshold of p<0.05 was utilized and the Tukey method was used for p value adjustment of estimated contrasts between each dose. Raw concentrations of each of the neurochemical analytes were log (base 10) transformed. For each experiment, the 11 samples collected from each rat at each of the three doses were grouped into 4 epochs, identified as ‘Pre-drug baseline’ (B1-B4), ‘drug’ (D1-D2), ‘post-drug’ (D3-D5), and ‘recovery’ (D6-D7), and averaged accordingly. A linear mixed model was implemented in which ‘dose’ and ‘state’ were treated as fixed factors and ‘subject’ was treated as a random intercept to account for intersubject variability. Separately for each dose, the model was used to compare the ‘drug,’ ‘post-drug,’ and ‘recovery’ epochs with the ‘pre-drug baseline’ epoch. To generate plots for neurochemical data, the ‘pre-drug baseline’ epoch was normalized to zero and changes relative to ‘pre-drug baseline’ were plotted for the ‘drug,’ ‘post-drug,’ and ‘recovery’ epochs. Mean values are shown for each dose with error bars indicating standard error of the mean (s.e.m.), unless noted otherwise. To assess the relationship between the levels of DMT, serotonin, and dopamine in the brain after DMT administration, Spearman’s rank correlation was used. Correlation coefficients (ρ) were calculated for each dose, and two-tailed t-tests with an alpha threshold of p<0.05 were conducted to test for statistical significance. For EEG analysis, a similar model was implemented, but each of the 7 drug epochs (D1-D7) were compared to the averaged ‘pre-drug baseline epoch’. Sex differences were tested with a separate linear mixed model in which ‘dose’, ‘state’, and ‘sex’ were treated as fixed factors and ‘subject’ was treated as a random intercept. The results of statistical comparisons between male and female rats have been provided in the **Extended Data Tables 5-1, 8-4, and 9-4**. An alpha threshold of p<0.05 was used as threshold for statistical significance and p values were adjusted for multiple comparisons via Dunnett’s test. Estimated contrasts between each state, as well as the percent change from baseline, t statistics, 95% confidence intervals (CI) and p values are reported for each comparison throughout the results section.

## Results

Histological analysis confirmed the location of OFM probes to be within mPFC (**Fig. 1C**) and S1BF (**Fig. 1D**).

### Effect of intravenous administration of DMT on behavior

The onset of behavioral effects typically began within 1 minute after the start of DMT infusion. The duration of behavioral changes showed a dose-dependent pattern with the effects lasting approximately 10-20 minutes after the low dose, 20-30 minutes after the medium dose, and 30-40 minutes after the high dose. For all doses, behavioral arousal was the first observed effect and occurred within the first 1-2 minutes after the start of the DMT infusion, and typically included exploratory behaviors such as rearing and sniffing. The majority of head twitches occurred during the 5-minute period immediately after the infusion of DMT. Average number of head twitches (±s.e.m.) during the 15-minute period starting at the beginning of the DMT infusions were: 6.6 ± 6.0 (low dose), 2.9 ± 2.1 (medium dose), and 2.6 ± 2.4 (high dose). The low dose caused significantly more head twitches when compared to the medium dose (t[31]=3.6, CI=0.34-6.78, p=0.28) and high dose (t[31]=4.0, CI=0.92-7.14, p=0.01) (**Fig. 2)**. Following the period of behavioral arousal and head twitches at the low dose, rats typically showed 10-20 minutes of inactivity and flattened body posture, with normal behavior returning approximately 30 minutes after the end of the infusion. During the 5-10 minute period after DMT infusion, the medium and high doses resulted in behaviors associated with serotonin syndrome, such as hind limb abduction and head weaving. The high dose showed the most pronounced manifestation of serotonin syndrome, which included all behaviors previously mentioned along with occasional forepaw treading, Straub tail, and backward walking. A period of inactivity followed these behaviors, which lasted for 20-30 minutes after the medium dose and 30-40 minutes following the high dose, after which the behavioral patterns returned to that observed during baseline wake state. There were no behavioral differences between males and females at any dose (**Fig. 2**).

**Figure 2.**
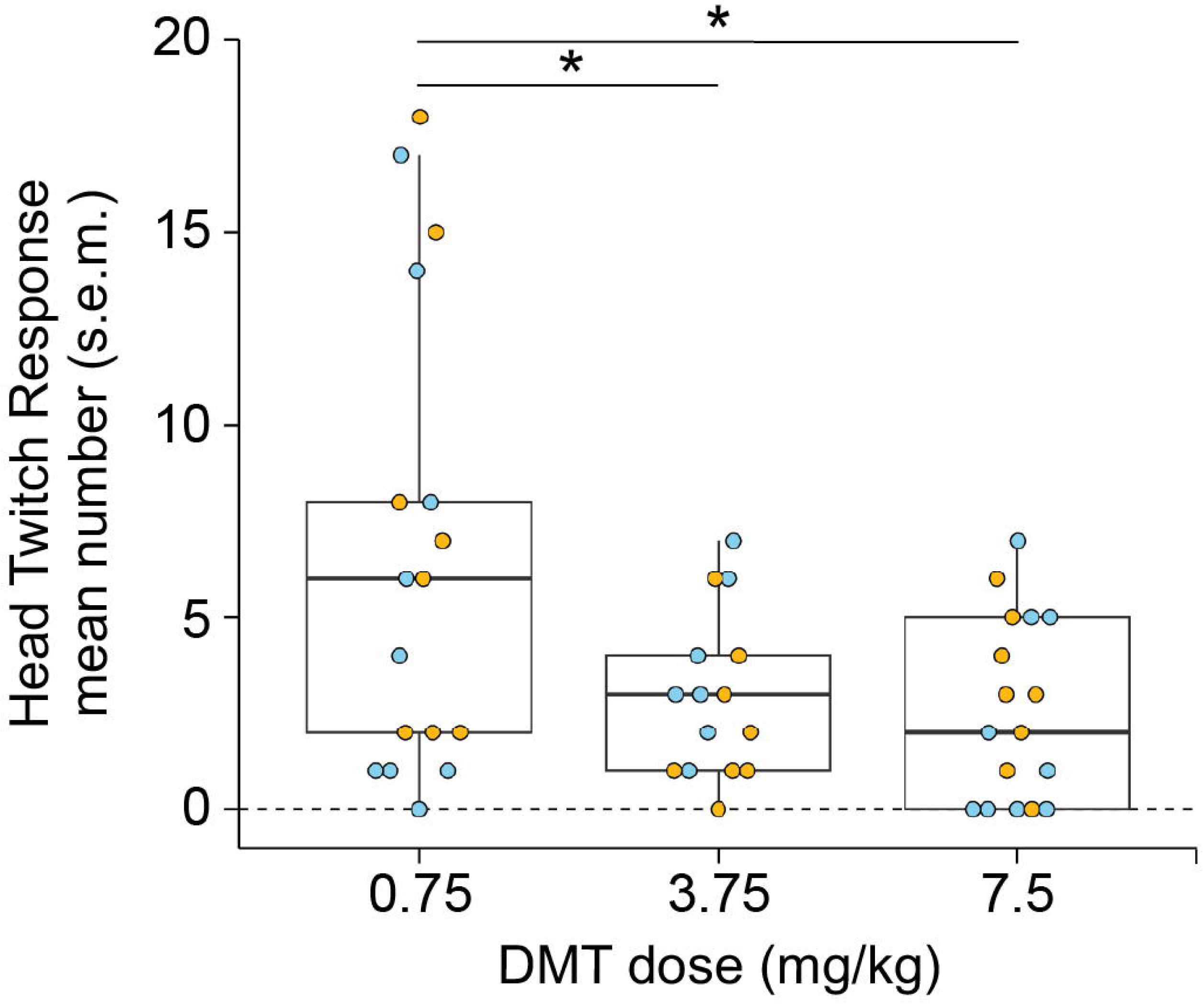
Head twitch response after intravenous infusion of DMT. As compared to the low dose (0.75 mg/kg), administration of DMT at medium (3.75 mg/kg) and high (7.5 mg/kg) doses resulted in significantly fewer head twitches. There were no differences in number of head twitches after any of the three doses between male and female rats. HTR was counted over a 15-minute window beginning at the start of DMT infusion. Individual data points are shown for each rat and colored by sex. Lines on box plots indicate median values, upper and lower bounds of the box indicate 75^th^ and 25^th^ quartiles, respectively, and whiskers indicate the range. A linear mixed model was used to compare the number of head twitches between doses of DMT. The p values were adjusted for multiple comparisons with Tukey’s test. *p<0.05.

### Effect of intravenous administration of DMT on cortical DMT concentration

DMT concentrations in mPFC and S1BF increased in a dose-dependent manner immediately after intravenous infusion of DMT (i.e. during the D1 epoch) (**Fig. 3**). The peak concentration (nM) in mPFC were (mean ± s.e.m.): 39.96 ± 7.76 for the low dose, 236.08 ± 60.03 for the medium dose, and 830.17 ± 405.09 for the high dose. The peak concentrations (nM) in S1BF were (mean ± s.e.m.): 38.14 ± 9.06 for the low dose, 183.11 ± 48.61 for the medium dose, and 369.70 ± 120.79 for the high dose. After the low-dose DMT infusion, the levels of DMT in mPFC (**Fig. 3A**) and S1BF (**Fig. 3B**) remained elevated until approximately 50 minutes after the start of DMT infusion (i.e., the D4 epoch). For the medium and high doses, the levels of DMT in both brain regions remained significantly higher than the baseline levels throughout the duration of the experiment (i.e., 87.5 minutes after the start of DMT infusion) (**Fig. 3A-B**). Statistical testing showed no effect of sex on the changes in cortical DMT levels following infusions. (**Extended Data Table 5-1**). The levels of DMT detected in mPFC and S1BF after intravenous infusion peaked during the D1 and D2 epochs and then gradually declined towards baseline. Therefore, to align the neurochemical analyses with the peak concentrations of DMT in the brain, we grouped the data into four states: pre-drug baseline (B1-B4), drug (D1-D2), post-drug (D3-D5), and recovery (D6-D7) (**Fig. 3C-D**).

**Figure 3.**
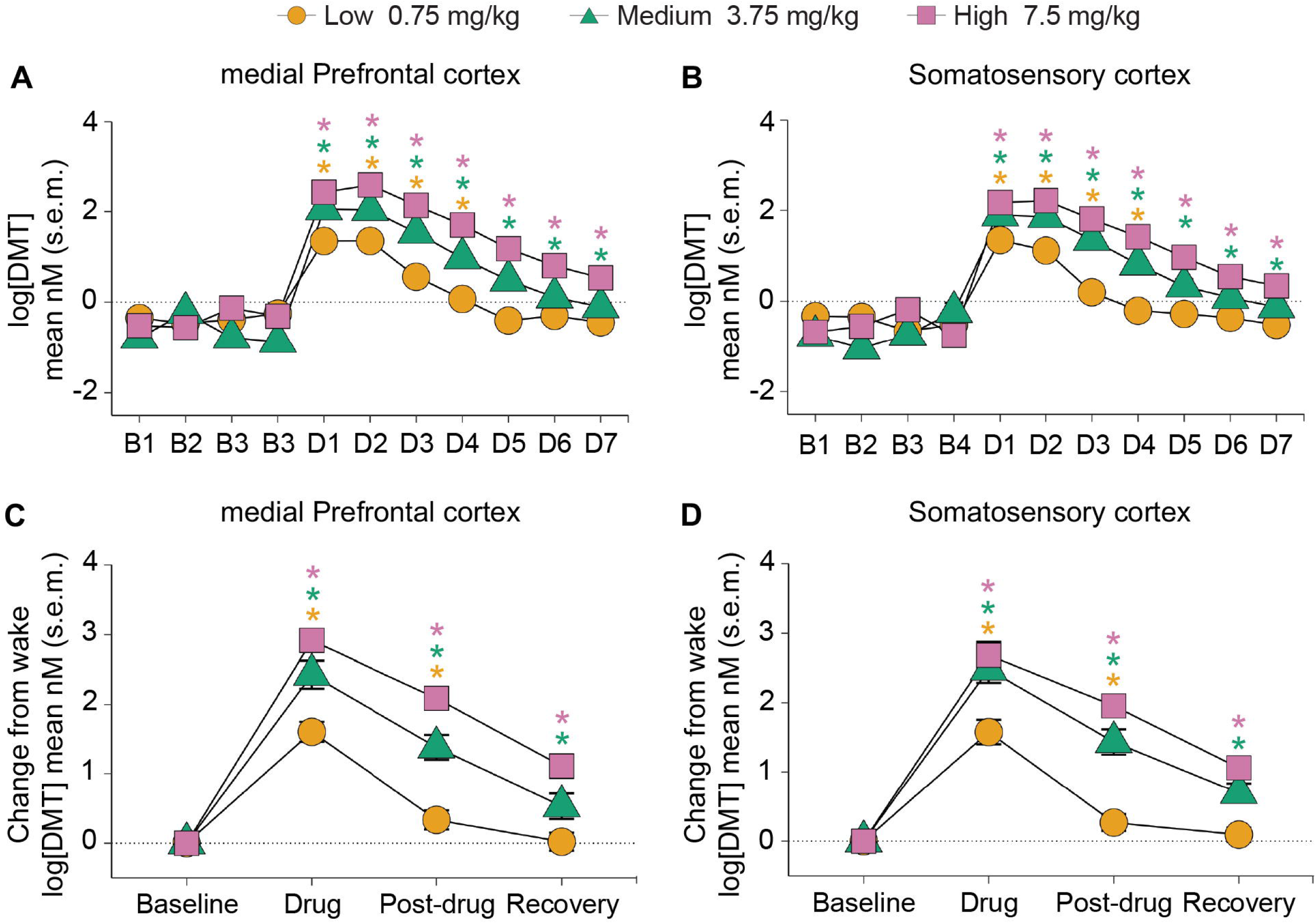
DMT levels in medial prefrontal cortex and somatosensory cortex before, during, and after intravenous administration of DMT. Panels (A) and (B) show log (base 10) transformed raw concentrations (nM) of DMT in medial prefrontal cortex (mPFC) and somatosensory cortex (S1BF) for each epoch, respectively. The pre-drug baseline data epochs are labelled as B1-B4. The data epochs collected during and after infusion of DMT are labelled as D1-D7. Each epoch shows open flow microperfusion sample collected over 12.5 minutes. Panels (C) and (D) show the log (base 10) transformed raw concentrations (nM) concentrations of DMT in mPFC and S1BF after combining the epochs B1-B4 as pre-drug “Baseline” (50 minutes), D1-D2 as “Drug” (25 minutes), D3-D5 as “Post-drug” (37.5 minutes), and D6-D7 as “Recovery” (25 minutes). The data are shown as change (log) from wake. Statistical analyses compared each epoch (D1-D7) in (A) and (B), and drug, post-drug, and recovery in (C) and (D) to the pre-drug baseline epoch. The DMT concentration remained significantly elevated until the D5 epoch for the low dose and remained significantly higher for medium and high doses throughout the duration of experiment. Orange circles represent the low dose (0.75 mg/kg), green triangles represent the medium dose (3.75 mg/kg), and magenta squares represent the high dose (7.5 mg/kg) DMT. A linear mixed model was used to compare the levels of recoverable DMT between epochs. The p values were adjusted for multiple comparisons via Dunnett’s test. *p<0.05 with asterisks color coded to match the symbol/dose. s.e.m.: standard error of the mean.

### Effect of intravenous administration of DMT on cortical neurochemistry

Intravenous DMT administration resulted in dose-dependent changes in the levels of 5-HT and dopamine in both mPFC and S1BF (**Fig. 4**). As compared to the pre-drug baseline state, intravenous administration of DMT at the medium and high doses caused a significant increase in 5-HT during the drug epoch in mPFC (medium dose: 221%, t[109]=5.73, p<0.001; high dose: 302%, t[109]=7.98, p<0.001) and S1BF (medium dose: 165%, t[116]=3.39, p=0.003; high dose: 184%, t[116]=3.85, p=0.001) (**Fig. 4A-B**). The low dose of DMT did not produce any statistically significant effect on 5-HT levels during the drug epoch but did cause a significant decrease in 5-HT concentration during the recovery period in both brain regions (mPFC: −66%, t[110]=-3.11, p=0.006; S1BF: - 66%, t[117]=-2.42, p=0.046) (**Fig. 4A-B**). Dopamine levels were affected only by the high dose of DMT, which showed a significant increase during the drug epoch in mPFC (178%, t[137]=4.51, p<0.001) and S1BF (146%, t[80]=2.77, p=0.02) (**Fig. 4C-D**). Apart from 5-HT and dopamine, we did not find any statistically significant changes during the drug epoch for the rest of the analytes but did observe changes during the post-drug and recovery epochs (**Fig. 5**). In mPFC, a significant decrease during both the post-drug and recovery epochs were observed for glycine (post-drug: −78%, t[153]=-.59, p=0.03; recovery: −78%, t[153]=-2.52, p=0.04), phenylalanine (post-drug: −79%, t[150]=-2.59, p=0.03; recovery: −77%, t[150]=-2.79, p=0.02), and taurine (post-drug: −81%, t[150]=-2.52, p=0.04; recovery: −80%, t[150]=-2.68, p=0.02); the decrease was observed only after administration of the high dose. The neurochemical changes in S1BF were as follows: low dose DMT resulted in a significant decrease during the recovery epoch in levels of glucose (−77%, t[141]=-3.22, p=0.005) and taurine (−85%, t[145]=-2.49, p=0.04). At the high dose, a significant decrease was observed during the recovery epoch in levels of adenosine (−76%, t[140]=-2.67, p=0.02) and glucose (−77%, t[141]=-3.10, p=0.007). The high dose also resulted in a significant decrease in levels of the following analytes during both the post-drug and recovery epochs: glycine (post-drug: −81%, t[145]=-3.33, p=0.003; recovery: −80%, t[145]=-3.55, p=0.001), phenylalanine (post-drug: −83%, t[141]=-3.12, p=0.006; recovery: −83%, t[141]=-3.0, p=0.009), and serine (post drug: −86%, t[145]=-2.43, p=0.04; Recovery:-85%, t145]=-2.58, p=0.03). Lastly, taurine levels were significantly reduced (−85%, t[145]=-2.49, p=0.04) during the post-drug epoch after the administration of high dose DMT. Basal concentrations of the 17 analytes from both mPFC and S1BF are provided in **Extended Data Table 5-1**. There were no differences observed between male and female rats in the changes in cortical neurochemistry following DMT infusions at any of the three doses. (**Extended Data Table 5-2**).

**Figure 4.**
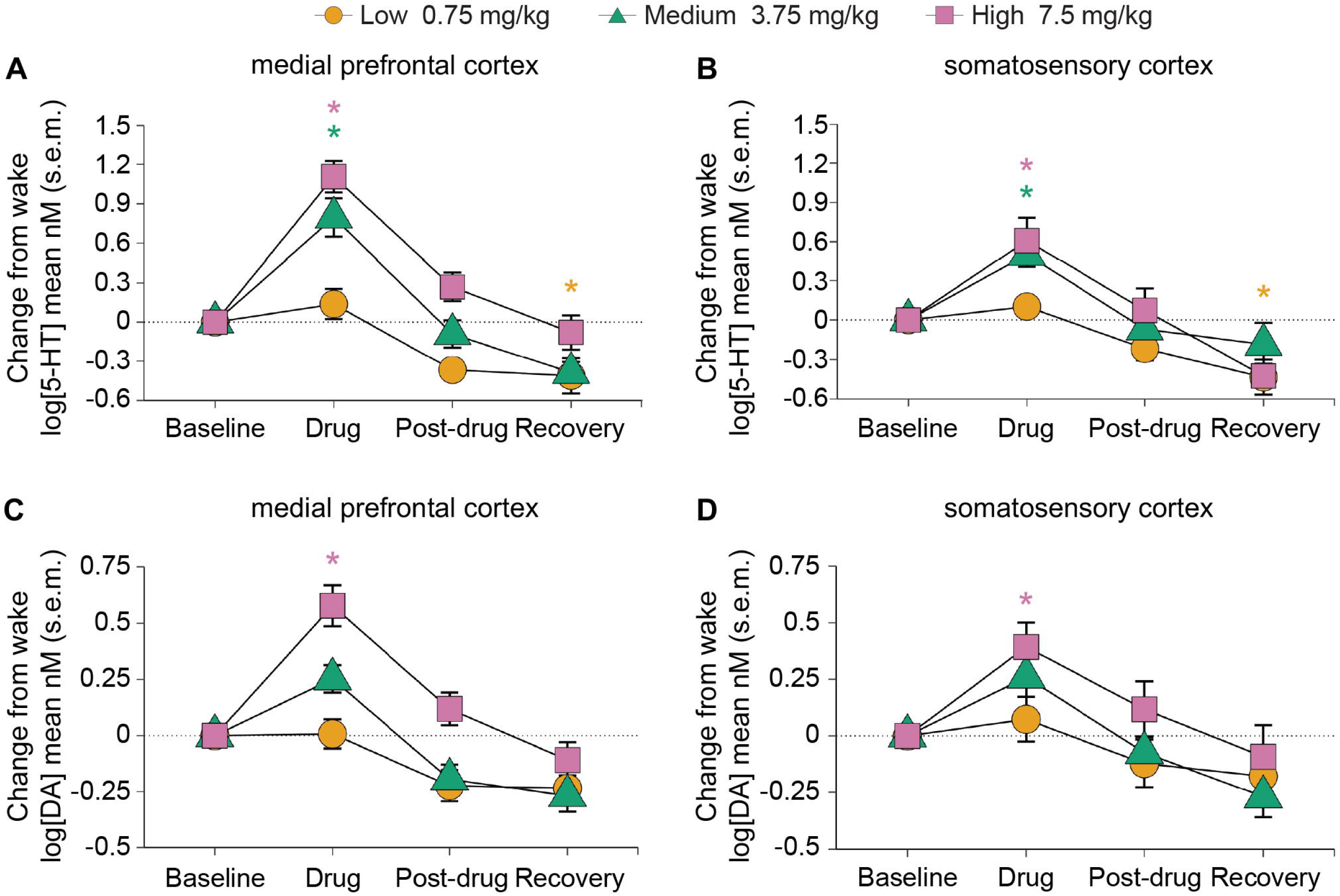
Intravenous administration of DMT increased the levels of 5-HT and dopamine in medial prefrontal cortex and somatosensory cortex. Panels (A) and (B) show log (base 10) transformed raw concentrations (nM) of 5-HT in medial prefrontal cortex (mPFC) and somatosensory cortex (S1BF), respectively. In both brain regions, medium and high doses of intravenous DMT caused 5-HT to increase during the drug epoch whereas the low dose DMT resulted in decreased 5-HT levels during the recovery epoch. Panels (C) and (D) show log (base 10) transformed raw concentrations (nM) of dopamine (DA) levels in mPFC and S1BF, respectively. The high dose of intravenous DMT caused DA to increase during the drug epoch in both brain regions. Orange circles represent the low dose (0.75 mg/kg), green triangles represent the medium dose (3.75 mg/kg), and magenta squares represent the high dose (7.5 mg/kg) DMT. A linear mixed model was used to compare the levels of 5-HT (A-B) or DA (C-D) between states. The p values were adjusted for multiple comparisons via Dunnett’s test. *p<0.05 with asterisks color coded to match the symbol/dose. s.e.m.: standard error of the mean.

**Figure 5.**
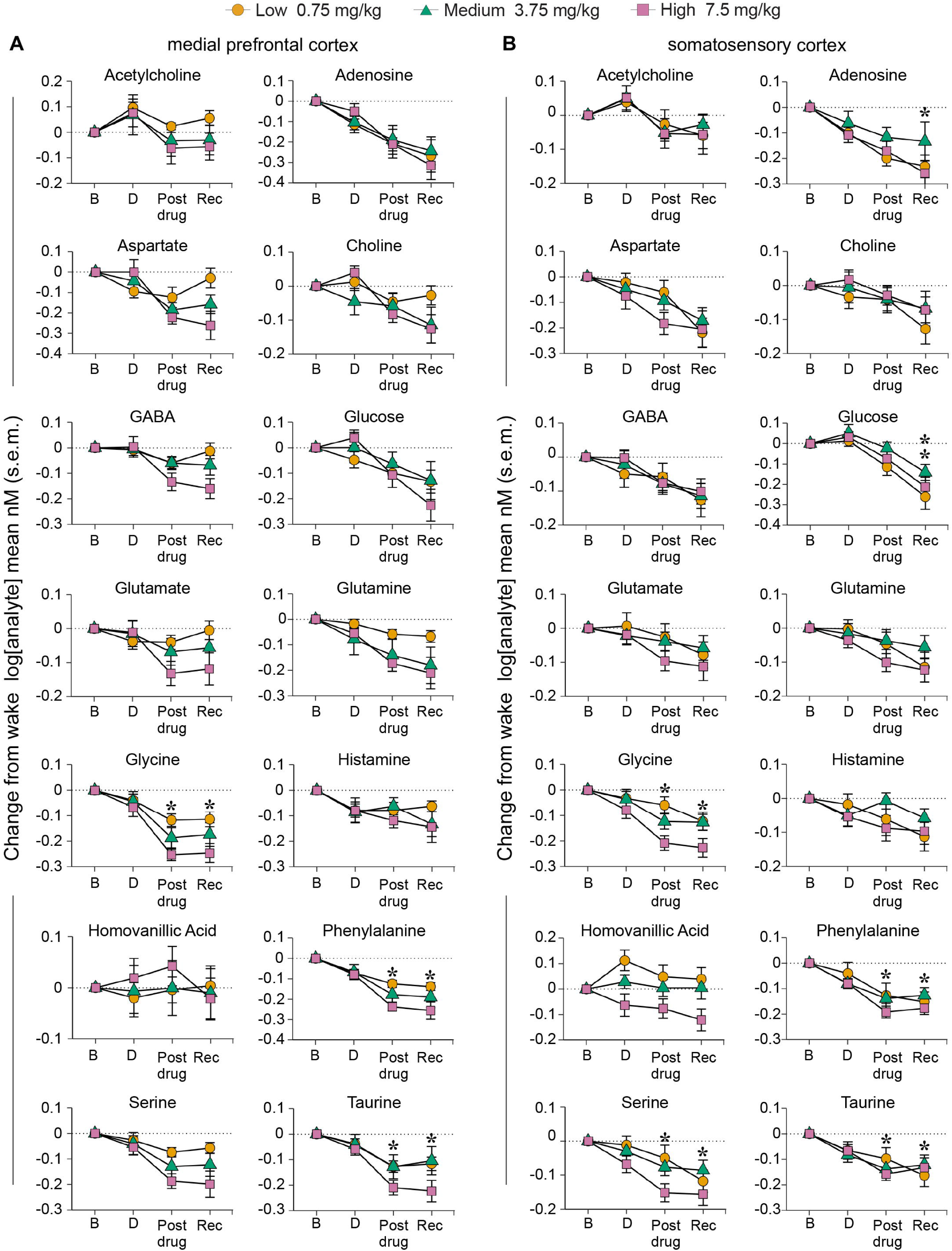
Neurochemical changes in medial prefrontal cortex and somatosensory cortex after intravenous administration of DMT. Log (base 10) transformed raw concentrations of neurochemicals quantified from medial prefrontal cortex (mPFC) and somatosensory cortex (S1BF) were averaged into pre-drug baseline (B), drug (D), post-drug (Post-drug), and recovery (Rec) bins. No significant changes, as compared to the wake epoch, were observed during the drug epoch for any of the analytes in either brain region. Orange circles represent the low dose (0.75 mg/kg), green triangles represent the medium dose (3.75 mg/kg), and magenta squares represent the high dose (7.5 mg/kg) DMT. A linear mixed model was used to compare levels of each analyte between states. The p values were adjusted for multiple comparisons via Dunnett’s test. *p<0.05 with asterisks color coded to match the symbol/dose. s.e.m.: standard error of the mean. See Extended Data Table 5-1 for a summary of baseline neurochemical concentrations in both brain regions, and Extended Data Table 5-2 for a summary of statistical comparisons of DMT-induced neurochemical changes.

Because the most salient neurochemical change following DMT administration was the increase in 5-HT and dopamine levels, we next assessed the relationship between the levels of 5-HT and dopamine and the associated levels of recoverable DMT in both brain regions during the D1-D7 epochs. Spearman correlation coefficients were computed using log transformed values of 5-HT, dopamine, and DMT in both mPFC and S1BF for each of the three doses. These analyses revealed significant positive correlation between 5-HT and recoverable DMT concentration as well as dopamine and recoverable DMT concentration, in both brain regions (**Fig. 6**). These correlations were largely dose-dependent, where the highest dose showed the strongest correlation for 5-HT in S1BF and for dopamine in both brain regions.

**Figure 6.**
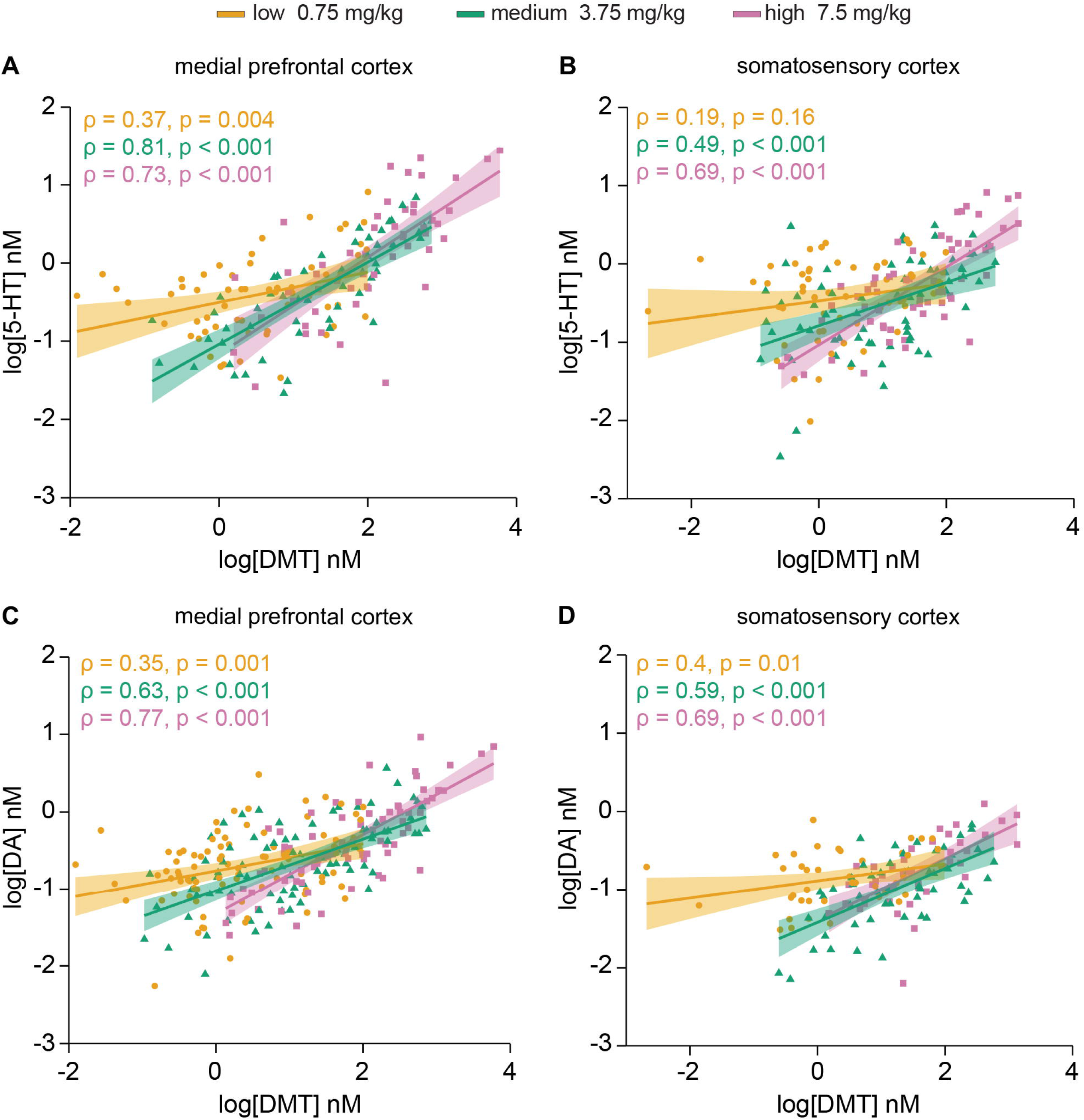
Relationship between the levels of recoverable DMT and levels of 5-HT and dopamine in medial prefrontal cortex and somatosensory cortex after intravenous administration of DMT. Correlation analyses using log (base 10) transformed neurochemical concentrations during D1-D7 epochs for low (0.75 mg/kg - orange), medium (3.75 mg/kg - green), and high (7.5 mg/kg – magenta) doses of DMT. Spearman’s rank correlation was used to calculate correlation coefficients (ρ) between serotonin (5-HT) and DMT levels in medial prefrontal cortex (mPFC) (A) and somatosensory cortex (S1BF) (B), and dopamine (DA) and DMT levels in mPFC (C) and S1BF (D). Regression lines are shown for each dose with shaded areas representing 95% confidence intervals. Positive correlations were observed between 5-HT and recoverable DMT and between DA and recoverable DMT in both brain regions. The p values were calculated via two-tailed t-tests. The correlation coefficients (ρ) are shown for each plot, with colors corresponding to dose.

Notably, endogenous DMT was detected during the pre-DMT baseline condition in 70% of the mPFC experiments and 80% of the S1BF experiments. The mean (± s.e.m.) DMT concentration was 0.66 ± 0.08 nM in mPFC and 0.54 ± 0.11 nM in S1BF, which was similar to that observed for 5-HT and dopamine in these brain regions (**Fig. 7**). **Extended Data Table 7-1** shows the concentrations of endogenous DMT detected in mPFC and S1BF during the pre-DMT baseline period in drug naïve condition (i.e., during baseline before the first experimental session) in each rat. We report that over 80% of the rats showed detectable levels of endogenous DMT in at least one brain region during vehicle infusions in the drug naïve.

**Figure 7.**
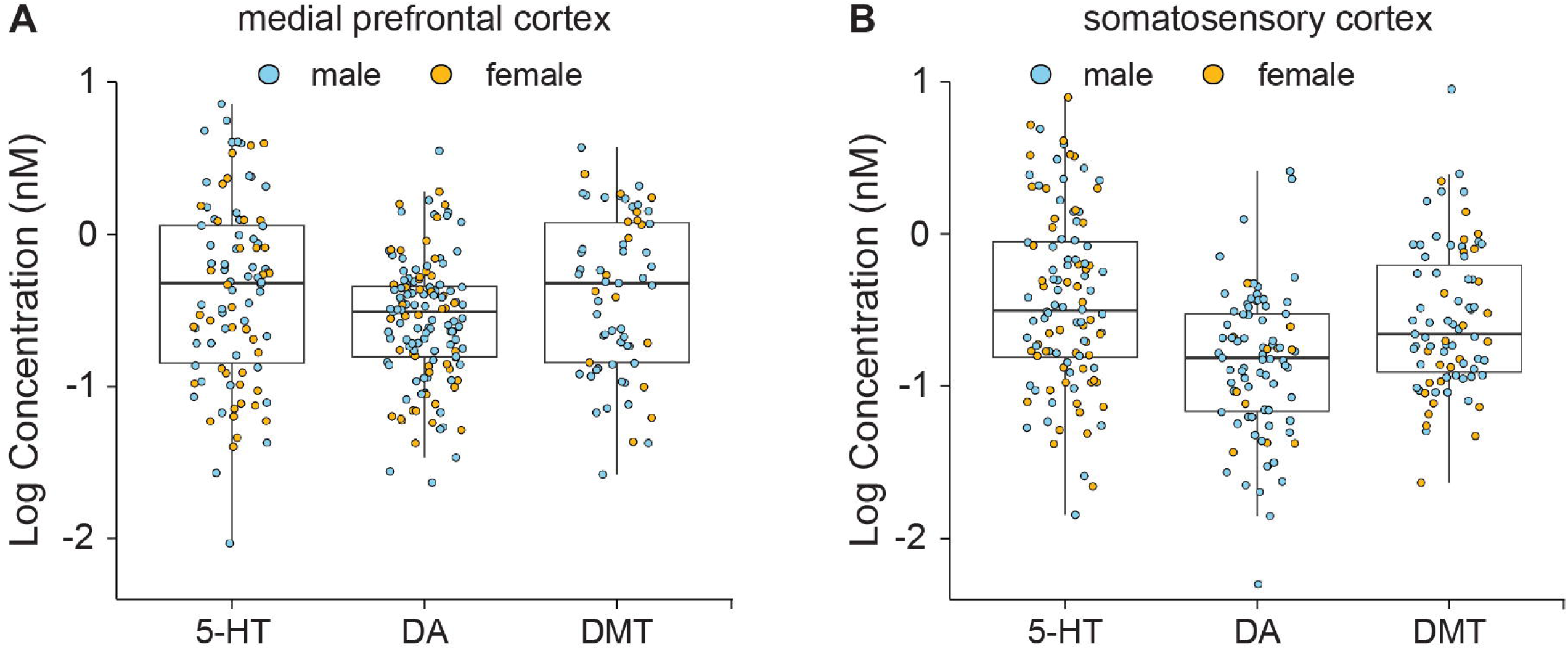
Basal levels of 5-HT, dopamine, and endogenous DMT in medial prefrontal cortex and somatosensory cortex. Basal values for 5-HT, dopamine, and DMT levels are shown for the four samples collected during the pre-drug baseline state across all experimental sessions. The levels of DMT are within the range of 5-HT and DA in both brain regions. Individual data points are shown for each rat and colored by sex. Lines on box plots indicate median values, upper and lower bounds of the box indicate 75^th^ and 25^th^ quartiles, respectively, and whiskers indicate the range. See Extended Data Table 7-1 for a summary of baseline endogenous DMT concentrations in drug naïve rats.

### Effect of intravenous administration of DMT on EEG spectral power

In addition to analyzing neurochemical changes in mPFC and S1BF, we simultaneously also conducted continuous high-density EEG recording to determine the effect of intravenous DMT on cortical neural dynamics. Statistically significant changes in EEG spectral power were observed after DMT infusions at each of the three doses (**Fig. 8**). As compared to the pre-drug baseline epoch, DMT infusion (i.e., D1 epoch) was characterized by a global increase in spectral power in delta band (low dose: 177%, t[519]=6.26, p<0.001; medium dose: 135%, t[519]=4.78, p<0.001) (**Fig. 8A**), medium gamma band (medium dose: ∼1%, t[519]=7.89, p<0.001; high dose: ∼1%, t[519]=5.18, p<0.001) (**Fig. 8D**), and high gamma band (low dose: ∼1%, t[519]=2.68, p=0.44; medium dose: ∼1%, t[519]=7.89, p<0.001; high dose: ∼1%, t[519]=4.74, p<0.001) (**Fig. 8E**). Simultaneously, there was a significant decrease in spectral power in theta band (low dose: −109%, t[519]=-8.07, p<0.001; medium dose: −100%, t[519]=-7.45, p<0.001) (**Fig. 8B**), and low gamma band (high dose: 1.1%, t[519]=-2.99, p=0.02) (**Fig. 8C**). Statistically significant changes were also observed during the D2-D7 epochs in delta, theta, and gamma bands at all three doses. The high dose DMT produced increase in delta power in D2 epoch, which remained elevated for the rest of the recording period, with statistically significant increase in D3 and D5 epochs. After the initial decrease in D1 epoch, the low gamma power showed a generalized increase at all doses for the rest of the recording period. Similarly, high dose DMT showed a delayed effect on theta power, which decreased significantly in D2 epoch, and then remained significantly low during the entire recoding period. Statistical testing showed no effect of sex on the changes in EEG spectral power between male and female rats at any dose. See **Extended Data Tables 8-1, 8-2, 8-3, and 8-4** for a summary of statistical comparisons of EEG power spectral changes.

**Figure 8.**
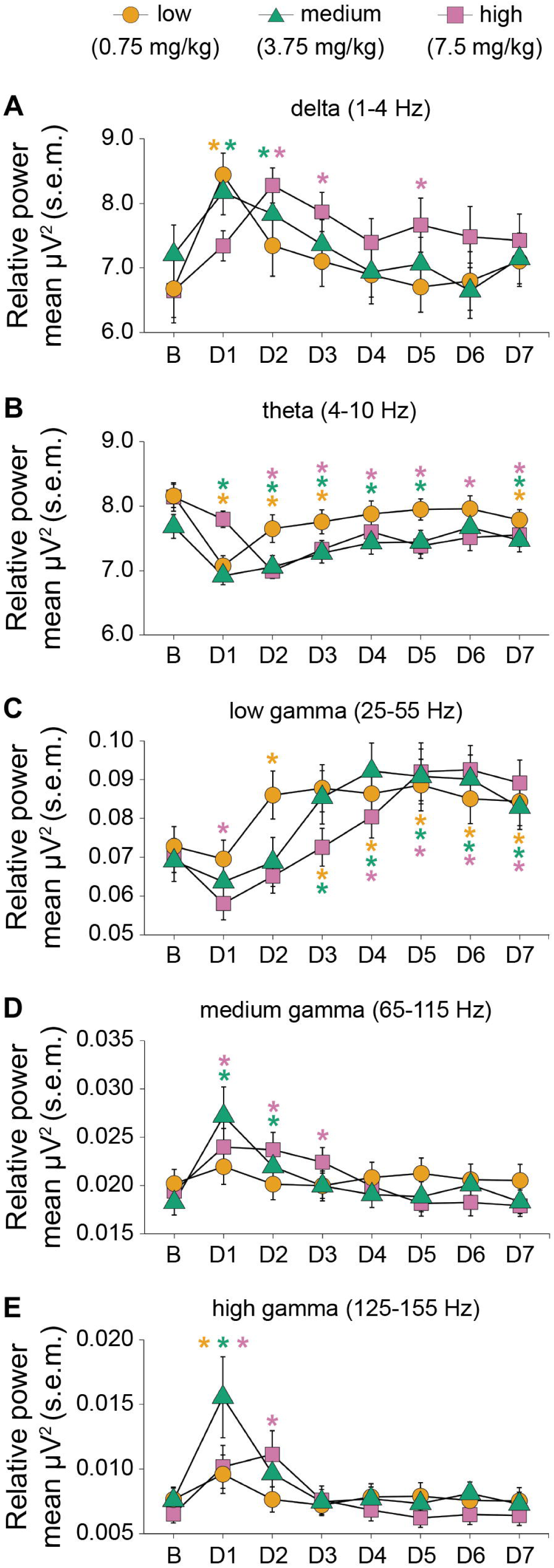
Effect of intravenous administration of DMT on EEG spectral power. The changes in EEG spectral power at selected frequency bands were determined by comparing the relative spectral power in the averaged pre-drug baseline epoch to that in each of the seven drug (D1-D7) epochs. There was increase in delta and decrease in theta relative power during the D1 epoch following the low dose (0.75mg/kg – Orange circles) and medium dose (3.75mg/kg – Green triangles) DMT, and during the D2 epoch following the high dose DMT (7.5mg/kg – Magenta squares). During D1, a significant increase in medium gamma power was observed after the medium and high doses of DMT, and increase in high gamma power after all three doses. Low gamma power increased in later post-infusion epochs (D4-D7) across doses. A linear mixed model was used to compare EEG relative power between states. The p values were adjusted for multiple comparisons via Dunnett’s test. *p<0.05 with asterisks color coded to match the symbol/dose. s.e.m.: standard error of the mean. See Extended Data Tables 8-1, 8-2, and 8-3 for a summary of statistical comparisons of changes in relative spectral power following DMT administration, and Extended Data Table 8-4 for a summary of statistical comparisons between male and female rats.

### Effect of intravenous administration of DMT on functional connectivity

As compared to the pre-drug baseline state, global delta connectivity was significantly reduced during the D1 epoch for the low dose (−15.4%, t[519]=-6.41, p<0.001) and medium dose (−9.7%, t[519]=-4.06, p<0.001) (**Fig. 9A**). For the high dose, this effect appeared after a delay, during the D2 epoch (− 16.8%, t[519]=-6.98, p<0.001). The high dose DMT also resulted in a significant increase in theta connectivity during the D1 epoch (5.7%, t[519]=5.96, p<0.001), followed by a significant decrease in theta connectivity during the D2 epoch (−3.7%, t[519]=-3.85, p=0.001) (**Fig. 9B**). There was a significant difference between males and females in theta connectivity (F[495]=2.49, p=.002). These differences were most apparent at the medium dose during the D1 epoch during which theta connectivity increased only in females (5.0%, t[495]=3.43, p=.004), and during the D2 epoch during which theta connectivity decreased only in males (−3.6%, t[495]=-3.0, p=0.02). At the high dose during the D2 epoch, theta connectivity decreased only for males (−3.4%, t[495]=-3.1, p=0.01). Finally, all three doses produced significant increase in connectivity during the D1 epoch across all three gamma bands: Low gamma band (low dose: 2.1%, t[519]=7.48, p<0.001; medium dose: 1.6%, t[519]=5.68, p<0.001; high dose: 4.9%, t[519]=9.19, p<0.001) (**Fig. 10A**), Medium gamma band (low dose: 1.0%, t[508]=3.55, p=0.003; medium dose: 1.7%, t[508]=5.88, p<0.001; high dose: 1.7%, t[508]=6.37, p<0.001) (**Fig. 10B**), and High gamma band (low dose: 3.3%, t[519]=6.25, p<0.001; medium dose: 5.9%, t[519]=11.02, p<0.001; high dose: 4.9%, t[519]=9.19, p<0.001) bands (**Fig. 10C**). A significant increase in global gamma connectivity persisted for both the medium and high doses and returned to baseline levels in a dose-dependent manner. Apart from theta connectivity, there were no significant sex differences in other connectivity measures. See **Extended Data Tables 9-1, 9-2, 9-3, and 9-4** for a summary of statistical comparisons of global connectivity changes.

**Figure 9.**
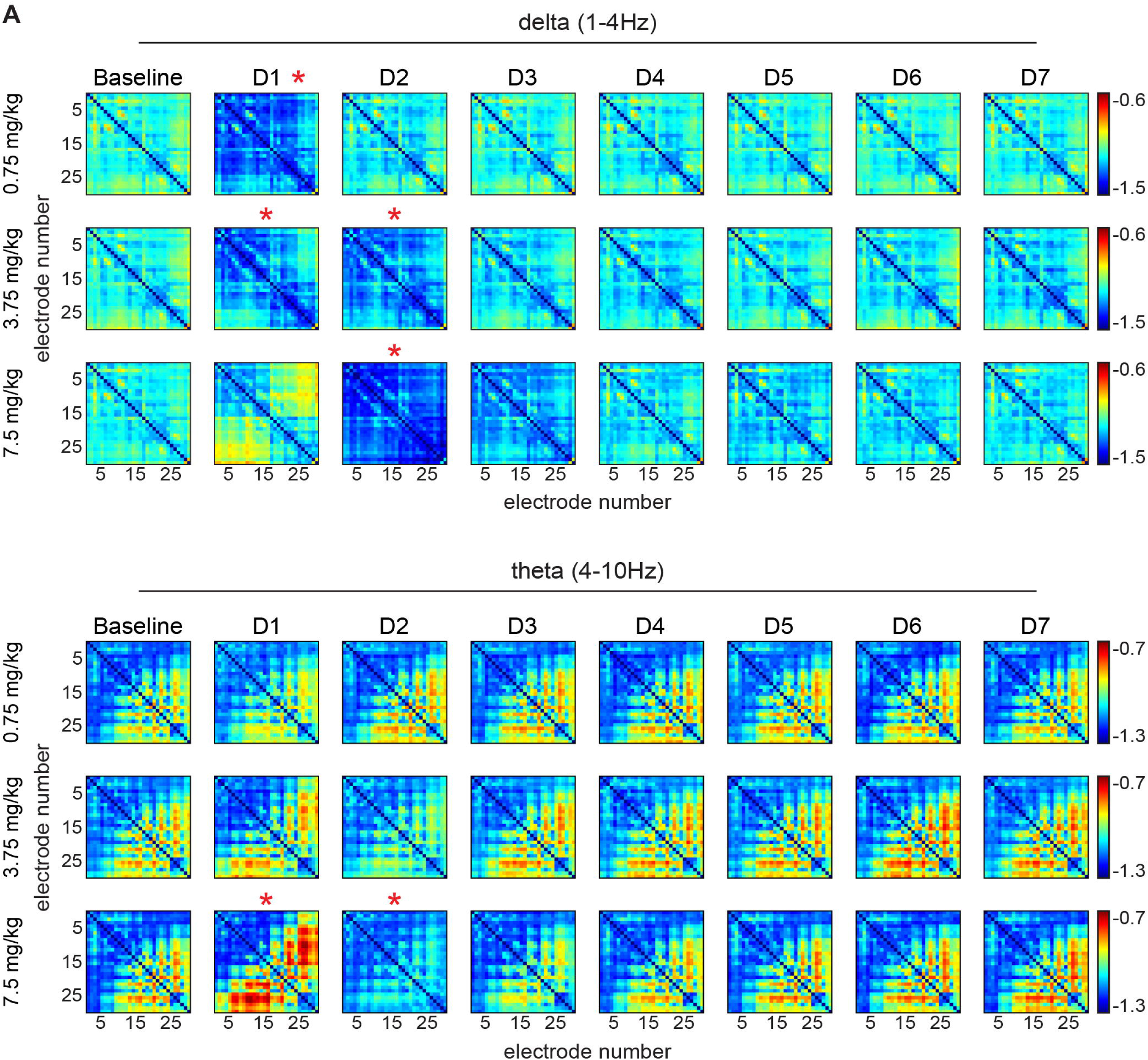
Effect of intravenous administration of DMT on delta- and theta functional connectivity. Adjacency matrices show the normalized weighted phase lag index (wPLI) value after low (0.75 mg/kg), medium (3.75 mg/kg), and high (7.5 mg/kg) doses of intravenous DMT for all channel pairs during pre-drug baseline state and each drug (D1-D7) epoch for delta (1-4 Hz) and theta (4-10 Hz) bands. Global delta wPLI decreased after the administration of low and medium doses of DMT during the D1 epoch, and after the high dose of DMT during the D2 epoch. Global theta wPLI increased after the high dose DMT during D1, primarily in frontal-posterior electrode pairs, and subsequently decreased during D2 epoch. Warmer colors indicate greater wPLI values (i.e., greater functional connectivity) and cooler colors indicate lower wPLI values. For electrode numbering schemes, see figure 1. A linear mixed model was used to compare global wPLI between states. The p values were adjusted for multiple comparisons via Dunnett’s test. *p<0.05. See Extended Data Tables 9-1, 9-2, and 9-3 for a summary of statistical comparisons of changes in wPLI following DMT administration, and Extended Data Table 9-4 for a summary of statistical comparisons between male and female rats.

**Figure 10.**
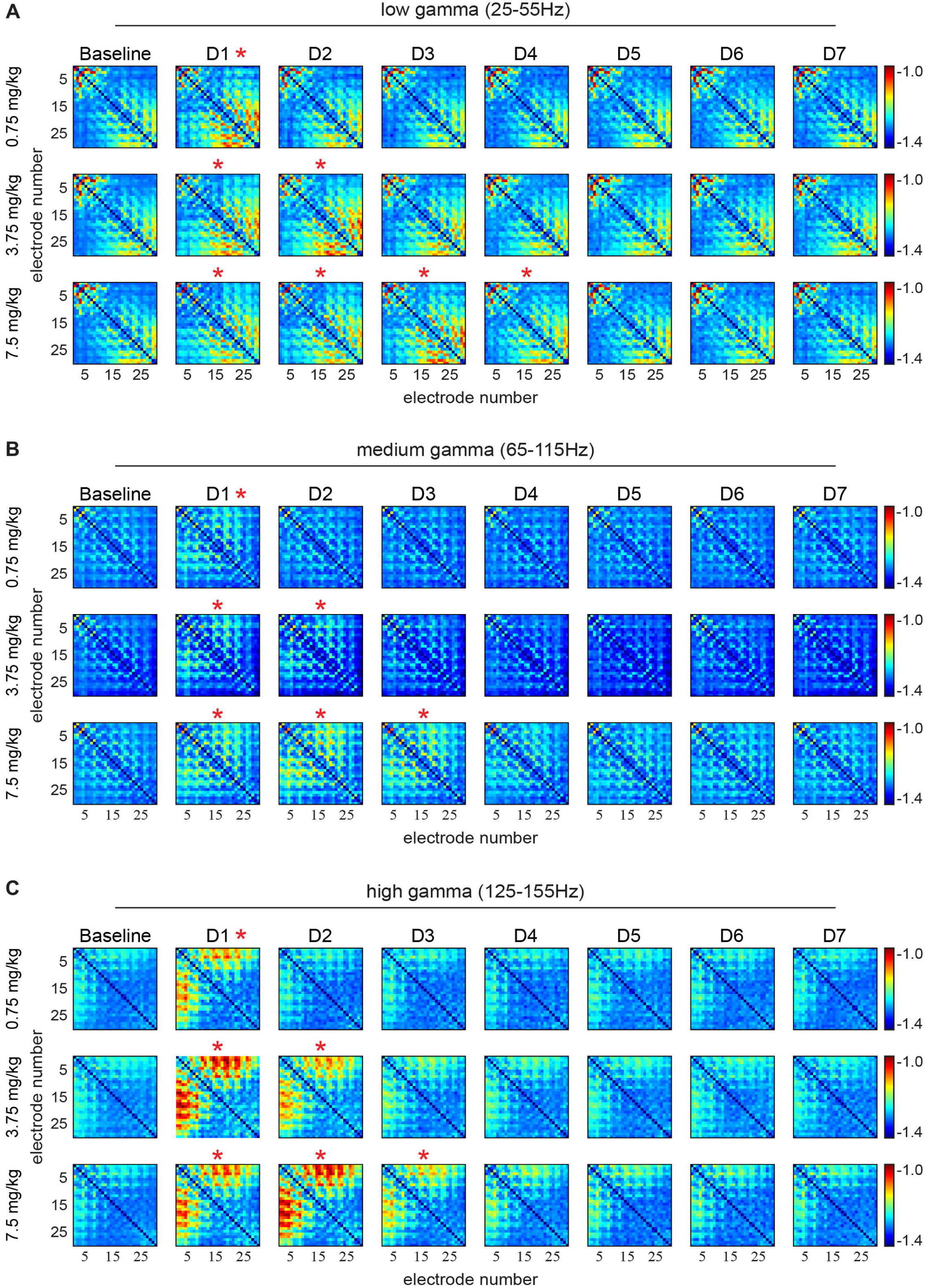
Effect of intravenous administration of DMT on gamma functional connectivity. Adjacency matrices show the normalized weighted phase lag index (wPLI) value after low (0.75 mg/kg), medium (3.75 mg/kg), and high (7.5 mg/kg) doses of intravenous DMT for all channel pairs during pre-drug baseline state and each drug (D1-D7) epoch for low (25–55 Hz), medium (65-115 Hz), and high (125-155 Hz) gamma bands. DMT administration increased functional connectivity in each of the three gamma bands in a dose-dependent manner, with higher doses causing more persistent changes, starting with the D1 epoch. Increases in low gamma connectivity dominated in parietal-occipital electrode pairs, medium gamma connectivity increases occurred in parietal electrodes, and high gamma connectivity increases were observed in frontal-parietal electrode pairs. Warmer colors indicate greater wPLI values (i.e., greater functional connectivity) and cooler colors indicate lower wPLI values. For electrode numbering schemes, see figure 1. A linear mixed model was used to compare global wPLI between states. The p values were adjusted for multiple comparisons via Dunnett’s test. *p<0.05

Of note, the global decrease in delta connectivity appears to be driven by frontal (no. 1-8) and parietal (no. 9-20) electrode pairs, as well as frontal-occipital (no. 21-30) electrode pairs (**Fig. 9A**). The changes in global theta and low gamma connectivity originated predominantly from parietal-occipital electrode pairs (**Fig. 9B, 10A**), whereas increase in medium gamma connectivity appears to be primarily driven by frontal-parietal electrode pairs (**Fig. 10B**). The changes in the high gamma connectivity (**Fig. 10C**) were primarily driven by the frontal electrode pairs.

## Discussion

In this study, we report that intravenous DMT caused a dose-dependent increase in 5-HT and dopamine in mPFC and S1BF, and produced head twitches — a behavioral surrogate for psychedelic drug action in rodents — which decreased with increasing doses. The most salient EEG changes were increase in spectral power and functional connectivity in the gamma bandwidth (See **Table 1** for a summary of DMT induced behavioral, neurochemical, and neurophysiological changes). Notably, we present the first report of endogenous DMT during normal wakefulness in prefrontal and somatosensory cortices at levels consistent with conventional neurotransmitters

Subcutaneous or intraperitoneal administration of DMT in rodents (Corne and Pickering, 1967; Stoff et al., 1978; Geyer et al., 1979; Jenner et al., 1980; Commissaris and Davis, 1983; González-Maeso et al., 2007; Cameron et al., 2018) has been reported to produce head twitches, hind limb abduction, low body posture and tremors, Straub tail, forepaw treading, and head weaving (Jenner et al., 1980). Our results are consistent with these previous findings and in addition demonstrated the dose-dependency of behavioral phenotypes. The incidence of head twitches was significantly higher following the low dose compared to medium or high doses. This pattern has been reported previously with DMT and other phenylalkylamine and tryptamine hallucinogens, where a substantial decrease in HTR occurs at higher doses (Fantegrossi et al., 2006, 2008; Halberstadt et al., 2020). This biphasic phenomenon has been posited to be mediated by differential receptor activation, with 5-HT_2A_ receptor activation driving the ascending portion of the HTR curve and 5-HT_2C_ receptor activation driving the descending portion (Fantegrossi et al., 2010). It is important to note that compared to the previous studies with serotonergic psychedelics, intravenous DMT administration in our study produced relatively low number of head-twitches, which could be due to at least two reasons. First, as opposed to other serotonergic psychedelics, DMT has a modest affinity at 5-HT2A receptors (Ki = 127 nM) (Keiser et al., 2009), which is the primary receptor that mediates head-twitch response. Second, few studies to date have investigated HTR in rats, and this is the first do so with intravenous administration, which allows rapid onset, but also rapid clearance from systemic circulation and the resulting faster offset of the effects. A previous study in mice (Cameron *et al.,* 2018) did not report head twitches after intraperitoneal administration of 10 mg/kg DMT, which is comparable to the high dose used in the current study.

Our rigorous technical approach to assessing neurochemical dynamics allowed for quantification of endogenous DMT during baseline normal wake condition in freely behaving rats, something that had only been done in a previous study of occipital cortex (Dean et al., 2019). We demonstrated that the levels of DMT in mPFC and S1BF during normal wakefulness are within the ranges of both 5-HT and dopamine, suggesting a physiological role for endogenous DMT, and supporting further investigation of DMT as a possible neurotransmitter. After intravenous administration, peak DMT levels in both cortical sites persisted for approximately 25 minutes and remained significantly elevated above baseline for more than 90 minutes for medium and high doses. This contrasts with the reported metabolic course of DMT in plasma after intravenous administration of hallucinogenic doses in humans; DMT concentrations peak at approximately 2 minutes and quickly decrease toward basal levels within 20 minutes (Timmermann et al., 2019). In rats, levels of recoverable DMT have been shown to be more than 5-fold higher in brain tissues as compared to that in blood, for up to 60 minutes after administration (Takahashi et al., 1985; Sitaram et al., 1987). These previous findings and the results reported in the current study suggest a potential storage or sequestration mechanism for DMT in the brain (Cozzi et al., 2009), or differential enzymatic clearance mechanisms between blood and brain tissues.

We found positive correlations between levels of recoverable DMT and levels of 5-HT and dopamine in mPFC and S1BF. Given the established role of 5-HT and dopamine in mediating mood and affect, these dose-dependent linear relationships suggest that DMT induced an altered state in rats, likely via an increase in endogenous neurotransmitters. Previous studies have shown that DMT acts as a 5-HT releaser, and has affinity at serotonin and dopamine transporters, which would presumably inhibit reuptake of the respective neurotransmitters, and cause increase in extracellular concentrations (Cozzi et al., 2009; Blough et al., 2014; Rickli et al., 2016). Additionally, DMT is not only a substrate, but also an inhibitor of monoamine oxidase (Ho et al., 1970; Ungar and Alivisatos, 1976; Barker et al., 1980), the primary enzyme responsible for metabolism of 5-HT and dopamine. Thus, sufficiently high concentrations of DMT may competitively inhibit monoamine oxidases, resulting in reduced metabolism of other endogenous monoamines. It has also been established that activation of 5-HT_2A_ receptors can directly regulate 5-HT levels in prefrontal cortex by increasing firing of dorsal raphe 5-HT neurons (Martı n-Ruiz et al., 2001) and dopamine levels through a feedback pathway involving efferent projections to the ventral tegmental area (Pehek et al., 2006; Alex and Pehek, 2007). We observed a significant decrease in phenylalanine — an essential amino acid precursor of tyrosine and hence catecholamine synthesis — in the post-drug and recovery period in both brain regions, which may indicate increased phenylalanine uptake into neurons to replenish depleted dopamine stores. Of note, a recent study (Wojtas et al., 2022) reported increase in 5-HT and dopamine levels in frontal cortex of rat after intraperitoneal administration of psilocybin and ketamine, which along with the results from the current study, suggests a shared mechanism of action of psychedelic drugs.

We also quantified the levels of several other neurotransmitters and signaling molecules, none of which showed a significant change during the acute drug state when DMT concentrations peaked in the brain. Of particular interest is the absence of a change in glutamate levels. This is unexpected considering the well-established finding that 5-HT_2A_ receptor activation causes increased glutamatergic excitatory postsynaptic currents in the mPFC, which is a region with high density of 5-HT_2A_ receptors (Aghajanian and Marek, 1999; Béïque et al., 2007). Accordingly, serotonergic psychedelics including LSD, psilocybin, and the 5-HT_2A_ agonist 2,5-dimethoxy-4-iodoamphetamine (DOI) have been reported to increase extracellular glutamate in the mPFC (Scruggs et al., 2003; Muschamp et al., 2004; Wojtas et al., 2022). Increased glutamatergic neurotransmission in layer V pyramidal neurons of the mPFC has been proposed as a key mechanism of psychedelic action and their neuroplasticity-promoting effects (Vollenweider and Kometer, 2010; Vollenweider and Preller, 2020). Although direct evidence for increased glutamate release in mPFC exists for other serotonergic psychedelics, this had previously not been established for DMT.

EEG results revealed persistent decreases in theta power at all three doses. This has been reported previously in human studies with psilocybin, LSD, and the atypical psychedelic, ketamine (Muthukumaraswamy et al., 2013; Pallavicini et al., 2019; Barnett et al., 2020). Previous human studies with DMT have also reported a reduction in alpha power as a neurophysiological correlate of DMT (Timmermann et al., 2019, 2023; Pallavicini et al., 2021). In contrast to humans where alpha oscillations are dominant, theta oscillations dominate in rats (Klimesch, 1999). Theta oscillations, which are generated in the hippocampus (Buzsáki, 2002), have long been associated with voluntary movement in rodents (Vanderwolf, 1969), but also play a major role in the synchronization of activity between hippocampal and cortical regions, which is required for learning, memory, and cognitive tasks (Zangbar et al., 2020). Thus, these results align with previous investigations and may be associated with disrupted communication between hippocampal and cortical regions following DMT.

Functional connectivity analyses revealed a pattern in which delta and theta connectivity diminished, giving way to increased gamma connectivity following DMT. For low and medium gamma frequencies, connectivity increases were observed primarily in electrode pairs involving parietal and occipital regions, whereas frontal electrode pairs were the primary contributors to the increased connectivity after the high gamma range. The duration and intensity of gamma changes were dose-dependent, with the highest dose causing the longest and most pronounced changes. Similar findings were reported in a recent rat study that demonstrated hypersynchronous high frequency gamma oscillations after serotonergic (LSD, DOI) and atypical psychedelics (ketamine) (Brys et al., 2023). In humans, changes in low gamma oscillations resulting from DMT (Pallavicini et al., 2021) and psilocybin (Kometer et al., 2015) correlate with several aspects of the subjective experience. Gamma rhythms represent fundamental components of cortical computation (Fries, 2009; Sohal et al., 2009) and are thought to play a major role in effective corticocortical connectivity (Fries, 2015). Therefore, such alterations in gamma oscillations may underlie the non-ordinary states of consciousness induced by psychedelic drugs.

Of note, to improve experimental rigor and translational applicability, we employed intravenous infusions of DMT, which allowed controlled and precise drug delivery into systemic circulation. In addition, we used OFM for neurochemical sampling, which offers two distinct advantages compared to conventional neurochemical sampling with microdialysis probes. First, unlike microdialysis probes, OFM probes do not extend beyond the guide tubes, thereby preventing damage to the blood-brain barrier and surrounding brain tissue, which eliminates potential confounds associated with inflammation and tissue scarring. Second, OFM probes do not use a semi-permeable membrane, which improves analyte recovery.

A limitation of our study is that the low temporal resolution of OFM sampling prevented assessment of transient or phasic neurochemical dynamics, potentially masking acute neurochemical changes. Furthermore, we did not investigate the effects of other psychedelic agents, which could have allowed a direct comparison to the current data, or serotonergic antagonists, which may have offered mechanistic insights into the action of DMT as it relates to neurochemical and neurophysiological changes reported in the current study. Antagonism studies have demonstrated that various 5-HT receptors contribute to behavioral, pharmacological, and neurophysiological outcomes of serotonergic psychedelics, suggesting that DMT may have similar effects (Erkizia-Santamaría; et al., 2022; Jaster et al., 2022; Tylš et al., 2023). Finally, the use of radiolabeled DMT could have provided definitive evidence to distinguish changes in endogenous DMT from levels of recoverable DMT resulting from drug administration. Despite these limitations, this study represents one of the most comprehensive neurochemical characterization of psychedelic drug action in rodents to date, and the neurophysiological changes mapped by functional connectivity suggests translational relevance of rodent models in investigating the neurobiology of DMT. Lastly, we demonstrated for the first time that DMT is present in prefrontal and somatosensory cortices of freely behaving normally behaving and awake rats at levels within the range of 5-HT and dopamine, a finding that motivates future investigation into the physiological role of DMT as a putative neurotransmitter.

## Acknowledgments

This research was supported by the National Institutes of Health Grant R01GM121919 to D.P., RF1-NS128522 to R.T.K., and funding from the Michigan Psychedelic Center and Department of Anesthesiology, University of Michigan Medical School, Ann Arbor, Michigan. The authors would like to thank Tiecheng Liu, M.D., Christopher Fields, B.S., and Lucy Wang, B.S., for assistance with experiments, and Chris Andrews, Ph.D. (*Consultant, Center for Consulting for Statistics, Computing & Analytics Research, University of Michigan*) for statistical consultation.

## Table Legends

**Table 1.** Summary of behavioral, neurophysiological, and neurochemical changes after intravenous administration of low (0.75 mg/kg), medium (3.75 mg/kg), and high (7.5 mg/kg) doses of DMT.

## Extended Data Legends

**Extended Data Table 1-1.** Multiple reaction monitoring conditions of 17 neurochemical analytes and their corresponding internal standards.

**Extended Data Table 5-1.** Basal concentrations of neurochemicals measured in medial prefrontal cortex (mPFC) and somatosensory cortex (S1BF).

**Extended Data Table 5-2.** Statistical comparison (ANOVA) of the DMT-induced neurochemical changes in medial prefrontal cortex (mPFC) and somatosensory barrel field cortex (S1BF) between male and female rats.

**Extended Data Table 7-1.** Endogenous concentrations of DMT in medial prefrontal cortex (mPFC) and somatosensory barrel field cortex (S1BF) in drug naïve rats.

**Extended Data Table 8-1.** Statistical comparisons of changes in relative spectral power after intravenous administration of low dose (0.75 mg/kg) DMT.

**Extended Data Table 8-2.** Statistical comparisons of changes in relative spectral power after intravenous administration of medium dose (3.75 mg/kg) DMT.

**Extended Data Table 8-3.** Statistical comparisons of changes in relative spectral power after intravenous administration of high dose (7.5 mg/kg) DMT.

**Extended Data Table 8-4.** Statistical comparison (ANOVA) of DMT-induced changes in relative spectral power between male and female rats

**Extended Data Table 9-1.** Statistical comparisons of changes in normalized weighted phase lag index (wPLI) after intravenous administration of low dose (0.75 mg/kg) DMT.

**Extended Data Table 9-2.** Statistical comparisons of changes in normalized weighted phase lag index (wPLI) after intravenous administration of medium dose (3.75 mg/kg) DMT.

**Extended Data Table 9-3.** Statistical comparisons of changes in normalized weighted phase lag index (wPLI) after intravenous administration of high dose (7.5 mg/kg) DMT.

**Extended Data Table 9-4.** Statistical comparison (ANOVA) of DMT-induced changes in normalized weighted phase lag index (wPLI) between male and female rats.

