## Extended Data Table 1-1 for "Neurochemical and Neurophysiological Effects of Intravenous Administration of *N,N*-Dimethyltryptamine in Rats"

**Extended Data Table 1-1.** Multiple reaction monitoring conditions of 17 neurochemical analytes and their corresponding internal standards.

| *Analyte* | *Precursor m/z* | *Product m/z* | *SRM Collision Energy* | *Retention Time (min)* | *Tube Lens* |
| --- | --- | --- | --- | --- | --- |
| Ch | 104 | 60 | 17 | 0.3 | 58 |
| d4-Ch | 108 | 60 | 17 | 0.3 | 58 |
| ACh | 146 | 87 | 13 | 0.4 | 53 |
| d4-ACh | 150 | 91 | 13 | 0.4 | 53 |
| ^13^C-Bz-Ser | 216.1 | 111 | 17 | 0.85 | 65 |
| Bz-Tau | 230 | 105 | 14 | 0.74 | 76 |
| ^13^C-Bz-Tau | 236 | 111 | 14 | 0.74 | 76 |
| Bz-Hist | 216 | 95 | 17 | 0.77 | 62 |
| ^13^C-Bz-Hist | 222 | 95 | 17 | 0.77 | 62 |
| Bz-Ser | 210 | 105 | 17 | 0.79 | 65 |
| Bz-Gln | 251 | 105 | 19 | 0.82 | 63 |
| ^13^C-Bz-Gln | 257 | 111 | 19 | 0.82 | 63 |
| Bz-Asp | 238 | 105 | 17 | 0.87 | 50 |
| ^13^C-Bz-Asp | 244 | 111 | 17 | 0.87 | 50 |
| Bz-Glc | 307 | 185 | 14 | 0.9 | 73 |
| ^13^C-Bz-Glc | 313 | 185 | 14 | 0.9 | 73 |
| Bz-Gly | 180 | 105 | 13 | 0.93 | 60 |
| ^13^C-Bz-Gly | 186 | 111 | 13 | 0.93 | 60 |
| DMT | 189.1 | 144.1 | 19 | 1 | 49 |
| d6-DMT | 195.1 | 144.1 | 19 | 1 | 64 |
| Bz-Glu | 252 | 105 | 15 | 0.96 | 69 |
| ^13^C-Bz-Glu | 258 | 111 | 15 | 0.96 | 69 |
| Bz-GABA | 208 | 105 | 14 | 1.15 | 61 |
| ^13^C-Bz-GABA | 214 | 111 | 14 | 1.15 | 61 |
| Bz-Ado | 372 | 136 | 27 | 1.4 | 76 |
| ^13^C-Bz-Ado | 378 | 136 | 27 | 1.4 | 76 |
| Bz-PA | 270 | 105 | 17 | 1.52 | 65 |
| ^13^C-Bz-PA | 276 | 111 | 17 | 1.52 | 65 |
| Bz-HVA | 304 | 105 | 16 | 1.64 | 66 |
| ^13^C-Bz-HVA | 310 | 111 | 16 | 1.64 | 66 |
| Bz-5-HT | 385 | 264 | 17 | 1.83 | 77 |
| ^13^C-Bz-5-HT | 397 | 270 | 17 | 1.83 | 77 |
| Bz-DA | 466 | 105 | 22 | 1.91 | 83 |
| ^13^C-Bz-DA | 484 | 111 | 22 | 1.91 | 83 |

ACh: Acetylcholine, Ado: Adenosine, Asp: Aspartate, Bz: Benzoylated, C_13_: Carbon-13, Ch: Choline, d: deuterium, DA: Dopamine, DMT: *N*,*N*-dimethyltrpytamine, GABA: γ-Aminobutryic acid, Glc: Glucose, Gln: Glutamine, Glu: Glutamate, Gly: Glycine, Hist: Histamine, HVA: Homovanillic acid, PA: Phenylalanine, Ser: Serine, Tau: Taurine.
