## Extended Data Table 5-1 for "Neurochemical and Neurophysiological Effects of Intravenous Administration of *N,N*-Dimethyltryptamine in Rats"

**Extended Table 5-1.** Basal concentrations of neurochemicals measured in medial prefrontal cortex (mPFC) and somatosensory barrel field cortex (S1BF).

|  |  | *Basal mPFC concentration* | | | |  | *Basal S1BF concentration* | | | |
| --- | --- | --- | --- | --- | --- | --- | --- | --- | --- | --- |
| *Analyte* |  | *n* | *Mean* | *SEM* | *Range* |  | *n* | *Mean* | *SEM* | *Range* |
| Acetylcholine |  | 141 | 1.80 | 0.16 | 0.28 – 13.57 |  | 143 | 2.82 | 0.28 | 0.18 – 18.69 |
| Adenosine |  | 167 | 13.76 | 1.15 | 0.38 – 96.55 |  | 172 | 10.29 | 0.62 | 0.62 – 47.84 |
| Aspartate |  | 141 | 1.80 | 0.16 | 0.15 – 547.20 |  | 174 | 71.77 | 64.60 | 51.27 – 62.11 |
| Choline |  | 173 | 238.03 | 19.62 | 32.34 – 1939.74 |  | 176 | 298.60 | 27.24 | 22.61 – 2434.54 |
| eDMT |  | 73 | 0.66 | 0.08 | 0.02 – 3.72 |  | 86 | 0.54 | 0.11 | 0.02 – 8.97 |
| Dopamine |  | 149 | 0.39 | 0.04 | 0.02 – 3.53 |  | 92 | 0.23 | 0.04 | 0.01 – 2.61 |
| GABA |  | 171 | 9.45 | 0.68 | 1.25 – 60.20 |  | 172 | 8.56 | 0.65 | 1.28 – 79.94 |
| Glucose^#^ |  | 179 | 248.74 | 14.71 | 7.29 – 1055.47 |  | 171 | 206.16 | 10.30 | 14.13 – 766.31 |
| Glutamate^#^ |  | 179 | 0.41 | 0.03 | 0.03 – 1.88 |  | 176 | 0.41 | 0.02 | 0.05 – 1.68 |
| Glutamine^#^ |  | 175 | 15.04 | 0.85 | 0.90 – 55.33 |  | 176 | 18.45 | 1.18 | 2.49 – 86.68 |
| Glycine^#^ |  | 180 | 2.16 | 0.16 | 0.31 – 10.85 |  | 176 | 2.66 | 0.14 | 0.30 – 11.48 |
| Histamine |  | 180 | 2.16 | 0.09 | 0.23 – 6.10 |  | 174 | 1.74 | 0.09 | 0.26 – 5.99 |
| HVA |  | 179 | 42.09 | 2.63 | 0.84 – 164.06 |  | 147 | 6.40 | 0.50 | 0.42 – 34.04 |
| Phenylalanine^#^ |  | 176 | 0.79 | 0.04 | 0.10 – 2.92 |  | 172 | 1.04 | 0.05 | 0.16 – 3.96 |
| Serine^#^ |  | 176 | 2.65 | 0.15 | 0.27 – 11.0 |  | 176 | 3.28 | 0.18 | 0.44 – 12.22 |
| 5-HT |  | 108 | 0.86 | 0.12 | 0.01 – 7.20 |  | 116 | 0.82 | 0.12 | 0.01 – 7.95 |
| Taurine^#^ |  | 176 | 1.59 | 0.09 | 0.21 – 6.68 |  | 176 | 1.69 | 0.08 | 0.17 – 6.62 |

‘n’ is the number of samples for each analyte and represents the sum of all available wake samples. eDMT: endogenous *N*,*N*-dimethyltryptamine, GABA: gamma-aminobutyric acid, HVA: Homovanillic acid, 5-HT: serotonin. #: µM concentration, all others expressed in nM.
