## Extended Data Table 5-2 for "Neurochemical and Neurophysiological Effects of Intravenous Administration of *N,N*-Dimethyltryptamine in Rats"

**Extended Table 5-2.** Statistical comparison (ANOVA) of the DMT-induced neurochemical changes in medial prefrontal cortex (mPFC) and somatosensory barrel field cortex (S1BF) between male and female rats.

|  |  | *mPFC* | | | *S1BF* | | | | |
| --- | --- | --- | --- | --- | --- | --- | --- | --- | --- |
| *Analyte* |  | *Sum of Squares* | *F statistic* | *P value* | |  | *Sum of Squares* | *F statistic* | *P value* |
| Acetylcholine |  | 0.133 | 1.043 | 0.386 | |  | 0.044 | 0.239 | 0.916 |
| Adenosine |  | 0.021 | 0.038 | 0.997 | |  | 0.387 | 1.431 | 0.224 |
| Aspartate |  | 0.701 | 1.169 | 0.325 | |  | 0.204 | 0.232 | 0.92 |
| Choline |  | 0.135 | 0.581 | 0.676 | |  | 0.051 | 0.229 | 0.922 |
| DMT |  | 0.794 | 0.842 | 0.5 | |  | 0.517 | 0.668 | 0.615 |
| Dopamine |  | 0.039 | 0.086 | 0.987 | |  | 0.279 | 1.298 | 0.276 |
| GABA |  | 0.143 | 0.568 | 0.686 | |  | 0.095 | 0.508 | 0.73 |
| Glucose |  | 0.312 | 1.019 | 0.398 | |  | 0.177 | 0.821 | 0.513 |
| Glutamate |  | 0.079 | 0.305 | 0.874 | |  | 0.091 | 0.708 | 0.587 |
| Glutamine |  | 0.074 | 0.329 | 0.858 | |  | 0.025 | 0.185 | 0.946 |
| Glycine |  | 0.167 | 0.611 | 0.655 | |  | 0.042 | 0.326 | 0.86 |
| Histamine |  | 0.005 | 0.024 | 0.999 | |  | 0.14 | 0.757 | 0.554 |
| HVA |  | 0.198 | 0.578 | 0.679 | |  | 0.164 | 0.713 | 0.584 |
| Phenylalanine |  | 0.039 | 0.216 | 0.929 | |  | 0.012 | 0.1 | 0.983 |
| Serine |  | 0.095 | 0.511 | 0.728 | |  | 0.039 | 0.299 | 0.878 |
| 5-HT |  | 0.583 | 0.924 | 0.451 | |  | 1.979 | 2.331 | 0.057 |
| Taurine |  | 0.122 | 0.68 | 0.606 | |  | 0.09 | 0.669 | 0.614 |

In addition to ‘dose’ and ‘state’, ‘sex’ was included as a fixed factor to test for potential differences in neurochemical responses to intravenous administration of DMT in male and female rats. ANOVA: Analysis of variance, DMT: *N*,*N*-dimethyltryptamine, GABA: gamma-aminobutyric acid, HVA: Homovanillic acid, 5-HT: serotonin.
