## Extended Data Table 7-1 for "Neurochemical and Neurophysiological Effects of Intravenous Administration of *N,N*-Dimethyltryptamine in Rats"

**Extended Table 7-1.** Endogenous concentrations of DMT in medial prefrontal cortex (mPFC) and somatosensory barrel field cortex (S1BF) in drug naïve rats.

|  | *DMT concentration (nM)* | |
| --- | --- | --- |
| *Rat ID* | *mPFC* | *S1BF* |
| *1* | nd | 0.24 |
| *2* | 0.18 | 0.27 |
| *3* | 0.88 | 0.12 |
| *4* | 0.22 | 0.13 |
| *5* | 0.12 | 0.10 |
| *6* | 0.80 | 0.31 |
| *7* | nd | nd |
| *8* | nd | nd |
| *9* | 1.16 | 0.25 |
| *10* | nd | 0.15 |
| *11* | 0.05 | 0.05 |
| *12* | 0.08 | 0.05 |
| *13* | nd | nd |
| *14* | 0.28 | 0.22 |
| *15* | 0.72 | 0.61 |
| *16* | 0.26 | 0.89 |
| *17* | nd | 0.12 |
| *18* | 0.67 | nd |
| *19* | 0.49 | nd |

The reported DMT levels are the average of the open flow microperfusion samples collected during the pre-drug baseline period. ‘nd’ indicates that DMT was not detected in experimental samples.
