## Extended Data Table 8-1 for "Neurochemical and Neurophysiological Effects of Intravenous Administration of *N,N*-Dimethyltryptamine in Rats"

**Extended Data Table 8-1.** Statistical comparisons of changes in relative spectral power after intravenous administration of low dose (0.75 mg/kg) DMT.

|  | |  | *Comparison* | *W ~ D1* | *W ~ D2* | *W ~ D3* | *W ~ D4* | *W ~ D5* | *W ~ D6* | *W ~ D7* |
| --- | --- | --- | --- | --- | --- | --- | --- | --- | --- | --- |
|  | ***δ*** | | Change | 1.772 | 0.673 | 0.431 | 0.220 | 0.032 | 0.125 | 0.434 |
|  |  |  | *t statistic* | 6.257 | 2.375 | 1.522 | 0.775 | 0.112 | 0.440 | 1.534 |
|  |  |  | p value | **<.001** | 0.098 | 0.484 | 0.904 | 1.000 | 0.983 | 0.476 |
|  | ***𝜃*** | | Change | -1.087 | -0.512 | -0.406 | -0.283 | -0.214 | -0.203 | -0.376 |
|  |  |  | *t statistic* | -8.070 | -3.800 | -3.010 | -2.103 | -1.589 | -1.507 | -2.791 |
|  |  |  | p value | **<.001** | **<.01** | **0.017** | 0.180 | 0.441 | 0.493 | **0.033** |
|  | ***⍺*** | | Change | -0.005 | <.01 | 0.039 | 0.021 | 0.028 | 0.025 | 0.024 |
|  |  |  | *t statistic* | -0.291 | 0.426 | 2.134 | 1.130 | 1.511 | 1.365 | 1.298 |
|  |  |  | p value | 0.995 | 0.984 | 0.168 | 0.733 | 0.491 | 0.586 | 0.629 |
|  | ***β*** | | Change | <.01 | 0.021 | 0.029 | 0.021 | 0.028 | 0.021 | 0.024 |
|  |  |  | *t statistic* | 0.962 | 2.444 | 3.367 | 2.450 | 3.300 | 2.478 | 2.829 |
|  |  |  | p value | 0.824 | 0.082 | **<.01** | 0.081 | **<.01** | 0.076 | **0.029** |
|  | ***Low 𝛾*** | | Change | -0.002 | 0.015 | 0.017 | 0.015 | 0.017 | 0.014 | 0.013 |
|  |  |  | *t statistic* | -0.450 | 4.096 | 4.589 | 4.197 | 4.815 | 3.835 | 3.643 |
|  |  |  | p value | 0.982 | **<.001** | **<.001** | **<.001** | **<.001** | **<.01** | **<.01** |
|  | ***Mid 𝛾*** | | Change | <.01 | <.001 | <.001 | <.01 | <.01 | <.01 | <.01 |
|  |  |  | *t statistic* | 2.061 | 0.290 | 0.137 | 0.961 | 1.379 | 0.739 | 0.662 |
|  |  |  | p value | 0.196 | 0.995 | 1.000 | 0.825 | 0.576 | 0.917 | 0.940 |
|  | ***High 𝛾*** | | Change | <.01 | <.001 | <.001 | <.001 | <.001 | <.001 | <.001 |
|  |  |  | *t statistic* | 2.684 | 0.224 | -0.325 | 0.466 | 0.540 | 0.128 | 0.073 |
|  |  |  | p value | **0.044** | 0.998 | 0.993 | 0.979 | 0.967 | 1.000 | 1.000 |

Estimated change in relative spectral power, comparing each of the drug conditions (D1 – D7) to the averaged pre-drug baseline value is shown. δ: 1–4 Hz, 𝜃: 4–10 Hz, ⍺: 10–15 Hz, β: 15–25 Hz, low 𝛾: 25–55 Hz, medium 𝛾: 65–115 Hz, high 𝛾: 125–155 Hz. Statistically significant changes where p<.05 are shown in bold.
