## Extended Data Table 8-2 for "Neurochemical and Neurophysiological Effects of Intravenous Administration of *N,N*-Dimethyltryptamine in Rats"

**Extended Data Table 8-2.** Statistical comparisons of changes in relative spectral power after intravenous administration of medium dose (3.75 mg/kg) DMT.

|  | *Comparison* | *W ~ D1* | *W ~ D2* | *W ~ D3* | *W ~ D4* | *W ~ D5* | *W ~ D6* | *W ~ D7* |
| --- | --- | --- | --- | --- | --- | --- | --- | --- |
| ***δ*** | Change | 1.352 | 1.007 | 0.539 | 0.111 | 0.236 | -0.178 | 0.321 |
|  | *t statistic* | 4.775 | 3.554 | 1.901 | 0.393 | 0.833 | -0.628 | 1.134 |
|  | p value | **<.001** | **<.01** | 0.266 | 0.988 | 0.882 | 0.949 | 0.730 |
| ***𝜃*** | Change | -1.003 | -0.867 | -0.653 | -0.492 | -0.482 | -0.254 | -0.454 |
|  | *t statistic* | -7.447 | -6.434 | -4.844 | -3.648 | -3.580 | -1.886 | -3.366 |
|  | p value | **<.001** | **<.001** | **<.001** | **<.01** | **<.01** | 0.273 | **<.01** |
| ***⍺*** | Change | -0.042 | 0.011 | 0.034 | 0.069 | 0.043 | 0.045 | 0.049 |
|  | *t statistic* | -2.288 | 0.598 | 1.842 | 3.715 | 2.351 | 2.417 | 2.663 |
|  | p value | 0.120 | 0.955 | 0.296 | **<.01** | 0.103 | 0.088 | **0.047** |
| ***β*** | Change | <.01 | 0.015 | 0.029 | 0.049 | 0.044 | 0.041 | 0.038 |
|  | *t statistic* | 0.199 | 1.757 | 3.455 | 5.824 | 5.159 | 4.825 | 4.446 |
|  | p value | 0.999 | 0.342 | **<.01** | **<.001** | **<.001** | **<.001** | **<.001** |
| ***Low 𝛾*** | Change | -0.004 | <.01 | 0.017 | 0.024 | 0.023 | 0.022 | 0.015 |
|  | *t statistic* | -1.241 | 0.170 | 4.767 | 6.649 | 6.263 | 6.075 | 4.057 |
|  | p value | 0.665 | 0.999 | **<.001** | **<.001** | **<.001** | **<.001** | **<.001** |
| ***Mid 𝛾*** | Change | <.01 | <.01 | <.01 | <.001 | <.001 | 0.001 | -0.001 |
|  | *t statistic* | 7.894 | 2.772 | 0.825 | -0.050 | -0.265 | 0.939 | -0.785 |
|  | p value | **<.001** | **0.034** | 0.885 | 1.000 | 0.997 | 0.835 | 0.901 |
| ***High 𝛾*** | Change | <.01 | <.01 | <.001 | <.001 | <.001 | <.001 | -0.001 |
|  | *t statistic* | 9.802 | 2.349 | -0.415 | -0.120 | -0.574 | 0.401 | -0.636 |
|  | p value | **<.001** | 0.104 | 0.986 | 1.000 | 0.961 | 0.987 | 0.946 |

Estimated change in relative spectral power, comparing each of the drug conditions (D1 – D7) to the averaged pre-drug baseline value is shown. δ: 1–4 Hz, 𝜃: 4–10 Hz, ⍺: 10–15 Hz, β: 15–25 Hz, low 𝛾: 25–55 Hz, medium 𝛾: 65–115 Hz, high 𝛾: 125–155 Hz. Statistically significant changes where p<.05 are shown in bold.
