## Extended Data Table 8-3 for "Neurochemical and Neurophysiological Effects of Intravenous Administration of *N,N*-Dimethyltryptamine in Rats"

**Extended Data Table 8-3.** Statistical comparisons of the changes in relative spectral power after intravenous administration of high dose (7.5 mg/kg) DMT.

|  |  | *Comparison* | *W ~ D1* | *W ~ D2* | *W ~ D3* | *W ~ D4* | *W ~ D5* | *W ~ D6* | *W ~ D7* |
| --- | --- | --- | --- | --- | --- | --- | --- | --- | --- |
| ***δ*** | | Change | 0.563 | 1.503 | 1.090 | 0.615 | 0.886 | 0.702 | 0.646 |
|  |  | *t statistic* | 1.988 | 5.307 | 3.850 | 2.171 | 3.129 | 2.480 | 2.282 |
|  |  | p value | 0.226 | **<.001** | **<.01** | 0.155 | 0.012 | 0.075 | 0.121 |
| ***𝜃*** | | Change | -0.321 | -1.123 | -0.789 | -0.515 | -0.733 | -0.602 | -0.563 |
|  |  | *t statistic* | -2.381 | -8.331 | -5.854 | -3.819 | -5.441 | -4.471 | -4.180 |
|  |  | p value | 0.096 | **<.001** | **<.001** | **<.01** | **<.001** | **<.001** | **<.001** |
| ***⍺*** | | Change | -0.049 | <.01 | -0.023 | -0.001 | <.01 | <.01 | 0.026 |
|  |  | *t statistic* | -2.661 | 0.204 | -1.234 | -0.064 | 0.484 | 0.476 | 1.432 |
|  |  | p value | 0.047 | 0.999 | 0.670 | 1.000 | 0.977 | 0.978 | 0.542 |
| ***β*** | | Change | -0.011 | 0.020 | <.01 | 0.018 | 0.040 | 0.038 | 0.037 |
|  |  | *t statistic* | -1.328 | 2.327 | 0.974 | 2.080 | 4.670 | 4.519 | 4.384 |
|  |  | p value | 0.610 | 0.109 | 0.818 | 0.188 | **<.001** | **<.001** | **<.001** |
| ***Low 𝛾*** | | Change | -0.011 | -0.004 | <.01 | 0.011 | 0.023 | 0.024 | 0.020 |
|  |  | *t statistic* | -2.992 | -1.047 | 1.005 | 3.168 | 6.385 | 6.530 | 5.587 |
|  |  | p value | **0.018** | 0.780 | 0.802 | **0.01** | **<.001** | **<.001** | **<.001** |
| ***Mid 𝛾*** | | Change | <.01 | <.01 | <.01 | <.01 | -0.001 | <.001 | -0.001 |
|  |  | *t statistic* | 5.183 | 4.904 | 3.648 | 1.219 | -0.524 | -0.427 | -0.763 |
|  |  | p value | **<.001** | **<.001** | **<.01** | 0.679 | 0.970 | 0.984 | 0.908 |
| ***High 𝛾*** | | Change | <.01 | <.01 | <.01 | <.001 | <.001 | <.001 | <.001 |
|  |  | *t statistic* | 4.737 | 5.943 | 1.489 | 0.522 | -0.247 | 0.083 | -0.006 |
|  |  | p value | **<.001** | **<.001** | 0.505 | 0.971 | 0.997 | 1.000 | 1.000 |

Estimated change in relative spectral power, comparing each of the drug conditions (D1 – D7) to the averaged pre-drug baseline value is shown. δ: 1–4 Hz, 𝜃: 4–10 Hz, ⍺: 10–15 Hz, β: 15–25 Hz, low 𝛾: 25–55 Hz, medium 𝛾: 65–115 Hz, high 𝛾: 125–155 Hz. Statistically significant changes where p<.05 are shown in bold.
