## Extended Data Table 8-4 for "Neurochemical and Neurophysiological Effects of Intravenous Administration of *N,N*-Dimethyltryptamine in Rats"

**Extended Table 8-4.** **Statistical comparison** (**ANOVA) of DMT-induced changes in relative spectral power between male and female rats**

| *Frequency band* |  | *Sum of Squares* | | *F statistic* | *P value* |
| --- | --- | --- | --- | --- | --- |
| Delta |  | 9.298 | 0.592 | | 0.872 |
| Theta |  | 2.611 | 0.726 | | 0.749 |
| Alpha |  | 0.075 | 1.2 | | 0.272 |
| Beta |  | 0.016 | 1.254 | | 0.233 |
| Low gamma |  | 0.003 | 1.064 | | 0.389 |
| Medium gamma |  | <.001 | 0.352 | | 0.986 |
| High gamma |  | <.001 | 1.412 | | 0.143 |

In addition to ‘dose’ and ‘state’, ‘sex’ was included as a fixed factor to test for potential sex differences in relative spectral power responses to intravenous DMT in male vs. female rats. ANOVA: Analysis of variance, δ: 1-4 Hz, 𝜃: 4-10 Hz, ⍺: 10-15 Hz, β: 15-25 Hz, low 𝛾: 25-55 Hz, medium 𝛾: 65-115 Hz, high 𝛾: 125-155 Hz. Statistically significant changes (p<0.05) are shown in bold.
