## Extended Data Table 9-1 for "Neurochemical and Neurophysiological Effects of Intravenous Administration of *N,N*-Dimethyltryptamine in Rats"

**Extended Data Table 9-1.** Statistical comparisons of changes in normalized weighted phase lag index (wPLI) after intravenous administration of low dose (0.75 mg/kg) DMT.

|  | |  | *Comparison* | *W ~ D1* | *W ~ D2* | *W ~ D3* | *W ~ D4* | *W ~ D5* | *W ~ D6* | *W ~ D7* |
| --- | --- | --- | --- | --- | --- | --- | --- | --- | --- | --- |
|  | ***δ*** | | Change | -0.154 | -0.033 | -0.011 | -0.029 | -0.016 | -0.012 | -0.011 |
|  |  |  | *t statistic* | -6.412 | -1.356 | -0.476 | -1.222 | -0.649 | -0.501 | -0.461 |
|  |  |  | p value | **<.001** | 0.591 | 0.978 | 0.677 | 0.943 | 0.974 | 0.980 |
|  | ***𝜃*** | | Change | 0.009 | 0.017 | 0.016 | 0.007 | 0.019 | 0.004 | 0.005 |
|  |  |  | *t statistic* | 0.975 | 1.811 | 1.712 | 0.782 | 1.983 | 0.385 | 0.542 |
|  |  |  | p value | 0.818 | 0.312 | 0.367 | 0.902 | 0.228 | 0.989 | 0.967 |
|  | ***⍺*** | | Change | 0.003 | 0.005 | 0.006 | 0.005 | <.001 | 0.005 | 0.005 |
|  |  |  | *t statistic* | 0.515 | 0.914 | 0.984 | 0.895 | -0.028 | 0.877 | 0.857 |
|  |  |  | p value | 0.972 | 0.847 | 0.813 | 0.856 | 1.000 | 0.864 | 0.872 |
|  | ***β*** | | Change | -0.011 | -0.003 | 0.003 | -0.001 | 0.004 | -0.003 | <.001 |
|  |  |  | *t statistic* | -2.379 | -0.623 | 0.598 | -0.191 | 0.768 | -0.546 | -0.078 |
|  |  |  | p value | 0.097 | 0.950 | 0.955 | 0.999 | 0.907 | 0.966 | 1.000 |
|  | ***Low 𝛾*** | | Change | 0.021 | 0.005 | -0.002 | -0.004 | 0.001 | <.001 | -0.001 |
|  |  |  | *t statistic* | 7.475 | 1.662 | -0.730 | -1.315 | 0.277 | -0.106 | -0.533 |
|  |  |  | p value | **<.001** | 0.396 | 0.919 | 0.618 | 0.996 | 1.000 | 0.969 |
|  | ***Mid 𝛾*** | | Change | 0.010 | -0.001 | -0.004 | -0.005 | -0.004 | -0.006 | -0.003 |
|  |  |  | *t statistic* | 3.548 | -0.540 | -1.553 | -1.907 | -1.499 | -2.177 | -1.019 |
|  |  |  | p value | **<.01** | 0.967 | 0.464 | 0.263 | 0.499 | 0.154 | 0.795 |
|  | ***High 𝛾*** | | Change | 0.033 | -0.001 | 0.001 | -0.001 | 0.003 | 0.001 | -0.002 |
|  |  |  | *t statistic* | 6.254 | -0.121 | 0.170 | -0.183 | 0.634 | 0.107 | -0.360 |
|  |  |  | p value | **<.001** | 1.000 | 0.999 | 0.999 | 0.947 | 1.000 | 0.991 |

Estimated change in connectivity, comparing each of the drug conditions (D1 – D7) to the averaged pre-drug baseline value is shown. δ: 1-4 Hz, 𝜃: 4-10 Hz, ⍺: 10-15 Hz, β: 15-25 Hz, low 𝛾: 25-55 Hz, medium 𝛾: 65-115 Hz, high 𝛾: 125-155 Hz. Statistically significant changes (p<0.05) are shown in bold.
