## Extended Data Table 9-2 for "Neurochemical and Neurophysiological Effects of Intravenous Administration of *N,N*-Dimethyltryptamine in Rats"

**Extended Data Table 9-2.** Statistical comparisons of changes in normalized weighted phase lag index (wPLI) after intravenous administration of medium dose (3.75 mg/kg) DMT.

|  | |  | *Comparison* | *W ~ D1* | *W ~ D2* | *W ~ D3* | *W ~ D4* | *W ~ D5* | *W ~ D6* | *W ~ D7* |
| --- | --- | --- | --- | --- | --- | --- | --- | --- | --- | --- |
|  | ***δ*** | | Change | -0.097 | -0.112 | -0.035 | -0.039 | -0.014 | -0.023 | -0.040 |
|  |  |  | *t statistic* | -4.059 | -4.678 | -1.473 | -1.626 | -0.589 | -0.944 | -1.645 |
|  |  |  | p value | **<.001** | **<.001** | 0.516 | 0.419 | 0.958 | 0.833 | 0.407 |
|  | ***𝜃*** | | Change | 0.005 | -0.018 | 0.012 | 0.010 | 0.009 | 0.015 | 0.013 |
|  |  |  | *t statistic* | 0.520 | -1.926 | 1.225 | 1.025 | 0.919 | 1.611 | 1.331 |
|  |  |  | p value | 0.971 | 0.254 | 0.675 | 0.792 | 0.845 | 0.427 | 0.607 |
|  | ***⍺*** | | Change | -0.018 | -0.005 | -0.001 | 0.008 | 0.003 | 0.002 | 0.009 |
|  |  |  | *t statistic* | -3.212 | -0.854 | -0.209 | 1.365 | 0.503 | 0.371 | 1.618 |
|  |  |  | p value | **<.01** | 0.874 | 0.998 | 0.585 | 0.974 | 0.990 | 0.423 |
|  | ***β*** | | Change | -0.005 | -0.001 | -0.009 | -0.002 | 0.002 | -0.004 | -0.002 |
|  |  |  | *t statistic* | -1.074 | -0.155 | -1.996 | -0.535 | 0.337 | -0.972 | -0.386 |
|  |  |  | p value | 0.765 | 0.999 | 0.222 | 0.968 | 0.993 | 0.819 | 0.989 |
|  | ***Low 𝛾*** | | Change | 0.016 | 0.021 | 0.007 | 0.004 | 0.002 | 0.001 | 0.003 |
|  |  |  | *t statistic* | 5.676 | 7.479 | 2.618 | 1.301 | 0.849 | 0.402 | 1.181 |
|  |  |  | p value | **<.001** | **<.001** | 0.053 | 0.627 | 0.875 | 0.987 | 0.702 |
|  | ***Mid 𝛾*** | | Change | 0.017 | 0.019 | 0.007 | 0.001 | -0.002 | -0.002 | 0.001 |
|  |  |  | *t statistic* | 5.875 | 6.713 | 2.640 | 0.414 | -0.694 | -0.588 | 0.485 |
|  |  |  | p value | **<.001** | **<.001** | 0.050 | 0.986 | 0.931 | 0.958 | 0.977 |
|  | ***High 𝛾*** | | Change | 0.059 | 0.037 | 0.008 | 0.004 | 0.000 | 0.003 | -0.003 |
|  |  |  | *t statistic* | 11.018 | 6.990 | 1.435 | 0.680 | -0.082 | 0.499 | -0.579 |
|  |  |  | p value | **<.001** | **<.001** | 0.540 | 0.935 | 1.000 | 0.974 | 0.960 |

Estimated change in connectivity, comparing each of the drug conditions (D1 – D7) to the averaged pre-drug baseline value is shown. δ: 1-4 Hz, 𝜃: 4-10 Hz, ⍺: 10-15 Hz, β: 15-25 Hz, low 𝛾: 25-55 Hz, medium 𝛾: 65-115 Hz, high 𝛾: 125-155 Hz. Statistically significant changes where p < .05 are shown in bold.
