## Extended Data Table 9-3 for "Neurochemical and Neurophysiological Effects of Intravenous Administration of *N,N*-Dimethyltryptamine in Rats"

**Extended Data Table 9-3.** Statistical comparisons of changes in normalized weighted phase lag index (wPLI) after intravenous administration of high dose (7.5 mg/kg) DMT.

|  | |  | *Comparison* | *W ~ D1* | *W ~ D2* | *W ~ D3* | *W ~ D4* | *W ~ D5* | *W ~ D6* | *W ~ D7* |
| --- | --- | --- | --- | --- | --- | --- | --- | --- | --- | --- |
|  | ***δ*** | | Change | 0.039 | -0.168 | -0.088 | -0.017 | -0.042 | -0.030 | -0.018 |
|  |  |  | *t statistic* | 1.610 | -6.976 | -3.655 | -0.707 | -1.747 | -1.248 | -0.743 |
|  |  |  | p value | 0.428 | **<.001** | **<.01** | 0.927 | 0.347 | 0.661 | 0.915 |
|  | ***𝜃*** | | Change | 0.057 | -0.037 | -0.009 | 0.006 | 0.009 | 0.010 | 0.015 |
|  |  |  | *t statistic* | 5.963 | -3.853 | -0.919 | 0.579 | 0.977 | 1.092 | 1.552 |
|  |  |  | p value | **<.001** | **0.001** | 0.845 | 0.960 | 0.817 | 0.755 | 0.464 |
|  | ***⍺*** | | Change | -0.010 | 0.008 | -0.001 | 0.002 | <.001 | 0.009 | 0.009 |
|  |  |  | *t statistic* | -1.780 | 1.388 | -0.260 | 0.298 | -0.078 | 1.564 | 1.574 |
|  |  |  | p value | 0.329 | 0.571 | 0.997 | 0.995 | 1.000 | 0.457 | 0.451 |
|  | ***β*** | | Change | -0.014 | 0.004 | -0.001 | -0.002 | -0.005 | -0.006 | -0.007 |
|  |  |  | *t statistic* | -3.089 | 0.871 | -0.223 | -0.520 | -1.108 | -1.343 | -1.428 |
|  |  |  | p value | **0.013** | 0.866 | 0.998 | 0.971 | 0.746 | 0.600 | 0.544 |
|  | ***Low 𝛾*** | | Change | 0.012 | 0.013 | 0.020 | 0.010 | 0.000 | 0.002 | 0.001 |
|  |  |  | *t statistic* | 4.410 | 4.677 | 7.400 | 3.592 | 0.091 | 0.562 | 0.200 |
|  |  |  | p value | **<.001** | **<.001** | **<.001** | **<.01** | 1.000 | 0.963 | 0.999 |
|  | ***Mid 𝛾*** | | Change | 0.017 | 0.025 | 0.020 | 0.006 | 0.005 | 0.003 | 0.000 |
|  |  |  | *t statistic* | 6.371 | 9.269 | 7.225 | 2.135 | 1.841 | 1.224 | 0.168 |
|  |  |  | p value | **<.001** | **<.001** | **<.001** | 0.168 | 0.296 | 0.676 | 0.999 |
|  | ***High 𝛾*** | | Change | 0.049 | 0.065 | 0.030 | 0.014 | 0.008 | 0.006 | 0.004 |
|  |  |  | *t statistic* | 9.191 | 12.252 | 5.614 | 2.580 | 1.586 | 1.152 | 0.748 |
|  |  |  | p value | **<.001** | **<.001** | **<.001** | 0.058 | 0.443 | 0.720 | 0.914 |

Estimated change in connectivity, comparing each of the drug conditions (D1 – D7) to the averaged wake value is shown. δ: 1-4 Hz, 𝜃: 4-10 Hz, ⍺: 10-15 Hz, β: 15-25 Hz, low 𝛾: 25-55 Hz, medium 𝛾: 65-115 Hz, high 𝛾: 125-155 Hz. Statistically significant changes (p<0.05) are shown in bold.
