## Extended Data Table 9-4 for "Neurochemical and Neurophysiological Effects of Intravenous Administration of *N,N*-Dimethyltryptamine in Rats"

**Extended Table 9-4.** Statistical comparison (ANOVA) of DMT-induced changes in normalized weighted phase lag index (wPLI) between male and female rats.

| *Frequency band* |  | *Sum of Squares* | | *F statistic* | *P value* |
| --- | --- | --- | --- | --- | --- |
| Delta |  | 0.035 | 0.324 | | 0.991 |
| Theta |  | 0.041 | 2.494 | | **0.002** |
| Alpha |  | 0.006 | 0.883 | | 0.578 |
| Beta |  | 0.003 | 0.764 | | 0.708 |
| Low gamma |  | 0.001 | 0.56 | | 0.896 |
| Medium gamma |  | <.001 | 0.346 | | 0.987 |
| High gamma |  | 0.003 | 0.533 | | 0.914 |

In addition to ‘dose’ and ‘state’, ‘sex’ was included as a fixed factor to test for potential differences in functional connectivity responses to intravenous DMT in male vs. female rats. δ: 1-4 Hz, 𝜃: 4-10 Hz, ⍺: 10-15 Hz, β: 15-25 Hz, low 𝛾: 25-55 Hz, medium 𝛾: 65-115 Hz, high 𝛾: 125-155 Hz. Statistically significant changes (p<0.05) are shown in bold.
