## Supplementary material for "Neurochemical and Neurophysiological Effects of Intravenous Administration of *N,N*-Dimethyltryptamine in Rats": Table 1

**Extended Dataset Tables**

Abbreviated title: Neural Correlates of DMT in Rats

Nicolas G. Glynos^1-4^, Emma R. Huels^2-5^, Amanda Nelson^2^, Youngsoo Kim^6^, Robert T. Kennedy^6^, George A. Mashour^2-5^, Dinesh Pal^1-5^**^*^**

1. Department of Molecular & Integrative Physiology, University of Michigan, Ann Arbor, MI 48109, USA
2. Department of Anesthesiology, University of Michigan, Ann Abor, MI 48109, USA
3. Michigan Psychedelic Center, University of Michigan, Ann Arbor, MI 48109, USA
4. Center for Consciousness Science, University of Michigan, Ann Arbor, MI 48109, USA
5. Neuroscience Graduate Program, University of Michigan, Ann Arbor, MI 48109, USA
6. Department of Chemistry, University of Michigan, Ann Arbor, MI 48109, USA

*****Correspondence:	Dinesh Pal, Ph.D.

7433 Medical Science Building 1

1150 West Medical Center Drive

Ann Arbor, Michigan 48109

Ph: 734-615-0234

**Extended Data Table 1-1.** Multiple reaction monitoring conditions of 17 neurochemical analytes and their corresponding internal standards.
